# Evaluating the epizootic and zoonotic threat of an H7N9 low pathogenicity avian influenza virus (LPAIV) variant associated with enhanced pathogenicity in turkeys

**DOI:** 10.1101/2024.04.16.589776

**Authors:** Joe James, Saumya S. Thomas, Amanda H. Seekings, Sahar Mahmood, Michael Kelly, Ashley C. Banyard, Alejandro Núñez, Sharon M. Brookes, Marek J. Slomka

**Affiliations:** Department of Virology, Animal and Plant Health Agency (APHA-Weybridge), Woodham Lane, Addlestone, Surrey KT15 3NB, UK; WOAH/FAO International Reference Laboratory for Avian Influenza, Animal and Plant Health Agency (APHA-Weybridge), Woodham Lane, Addlestone, Surrey KT15 3NB, UK; Pathology and Animal Sciences Department, Animal and Plant Health Agency (APHA-Weybridge), Woodham Lane, Addlestone, Surrey KT15 3NB, UK

**Author notes:** Authors contributed equally, order is alphabetical (Joe James, Saumya S. Thomas).

**Keywords:** avian influenza virus, H7N9, avian, transmission, pathogenicity, host adaptation, L226Q

## Abstract

Between 2013-2017, A/Anhui/1/13-lineage (H7N9) low pathogenicity avian influenza virus (LPAIV) was epizootic in chickens in China causing mild disease, with 616 fatal human cases. Despite poultry vaccination, H7N9 has not been eradicated. Previously we demonstrated increased pathogenesis in turkeys infected with H7N9, correlating with the emergence of the L217Q (L226Q H3 numbering) polymorphism in the haemagglutinin (HA) protein. A Q217 containing virus also arose and is now dominant in China following vaccination. We compared infection and transmission of this Q217 containing ‘turkey-adapted’ (ty-ad) isolate alongside the H7N9 (L217) *wild-type* (*wt)* virus in different poultry species, and investigated the zoonotic potential in the ferret model. Both *wt* and ty-ad viruses demonstrated similar shedding and transmission in turkeys and chickens. However, the ty-ad virus was significantly more pathogenic than the *wt* virus in turkeys but not in chickens, causing 100% and 33% mortality in turkeys respectively. Expanded tissue tropism was seen for the ty-ad virus in turkeys but not chickens, yet the viral cell receptor distribution was broadly similar in visceral organs of both species. The ty-ad virus required exogenous trypsin for *in vitro* replication yet had increased replication in primary avian cells. Replication was comparable in mammalian cells and the ty-ad virus replicated successfully in ferrets. The L217Q polymorphism also affected antigenicity. Therefore, H7N9 infection in turkeys can generate novel variants with increased risk through altered pathogenicity and potential HA antigenic escape. These findings emphasise the requirement for enhanced surveillance and understanding of A/Anhui/1/13-lineage viruses and their risk to different species.

## INTRODUCTION

In 2013 the first human infection with a novel reassortant H7N9 avian-origin influenza A virus (IAV) was reported in China [1]. To date, there have been 1568 laboratory-confirmed human cases of H7N9 which include 616 fatal cases, giving a case fatality rate of 39% [2, 3], many of which have been associated with human exposure to infected poultry, predominantly at live bird markets [4]. While chickens, pigeons, and ducks have all been reported as positive for H7N9, chickens are considered as the maintenance host [5, 6]. H7N9 achieved enzootic levels in chickens across large parts of China, circulating in the absence of clinical disease as a low pathogenicity avian influenza virus (LPAIV) [5, 6]. After almost four years of extensive circulation in China, a high pathogenicity AIV (HPAIV) H7N9 variant was detected among one of the two major LPAIV H7N9 clades which had evolved during this period [7, 8]. Vaccination of poultry against H7N9 commenced in China in 2017 during the fifth wave and achieved apparent success in reducing detections of human infections [9]. However, despite the reduction of zoonotic H7N9 cases [9], this virus has not been eradicated from poultry in China, and sporadic detection in poultry and the environment of farms and live bird markets have continued across different provinces of China (Fujian, Guangdong and Henan) [3], suggesting continued circulation in Chinese poultry. Genetic analysis of H7N9 viruses which continue to circulate following the adoption of vaccination have consistently identified stable haemagglutinin (HA) gene polymorphisms, namely the change from a Leucine (L) to Glutamine (Q) amino acid residue at position 217 in the H7 mature peptide (L217Q: L226Q H3 numbering; L235Q complete protein numbering)) [10, 11].

Amino acid position 217 is located in the 220-loop of the HA protein forming part of the influenza A virus receptor binding site. Possession of a L at position 217 has been previously shown to increase the affinity of HA binding to ‘human-like’, α-2,6 sialic acid (SA), while a Q at position 217 increases affinity of binding to ‘avian-like’, α2,3 SA [12–14]. Thus, the Q217L polymorphism present in wave 1-5 (2013-2017) H7N9 viruses, along with other molecular motifs such E627K in the basic polymerase 2 (PB2) protein, have been attributed to the zoonotic potential of these viruses [15, 16]. These human H7N9 cases have followed a seasonal pattern with five successive winter waves [17]. The fifth wave of H7N9 (2016-2017) attained its widest geographical distribution in China [17], reinforcing earlier concerns about the spread of H7N9 to neighbouring countries [18]. Travel-related human cases have resulted in the detection of H7N9 in Canada and Malaysia [19, 20], while confiscated poultry meat produce smuggled from China to Japan was found to be contain H7N9 LPAIV and HPAIV [21]. In addition, H7N9 has been demonstrated to be able to reassort with co-circulating H9N2 viruses, producing novel viruses with altered biological properties [22, 23]. Such events may increase the risk of emergence of infection in non-chicken hosts, including other avian species, although the requirement and role of further adaptation to different bird species is unclear. Experimental investigations of H7N9 LPAIV in poultry have focused almost exclusively on chickens [24]. Turkeys represent a major poultry species and important food source globally[25], although the turkey sector in China is comparatively low compared to duck and chicken sectors [25]. In addition, turkeys have been reported as being highly susceptible to a range of different AIV strains, including H7 subtypes [26, 27], and both wild and domestic turkeys have been highly affected during the current H5N1 clade 2.3.4.4b H5N1 panzootic [28, 29]. We previously investigated the effects of H7N9 infection and transmission in turkeys [24] and, using the prototypical wild-type (*wt*) H7N9 LPAIV of human origin (A/Anhui/1/2013), we demonstrated that direct-infection of turkeys resulted in onward transmission to contact turkeys. Moreover, we observed an unexpectedly dramatic increase in pathogenicity for a LPAIV, resulting in clinical disease and significant turkey mortality [24]. Disease severity correlated with the rapid and consistent emergence of the L217Q polymorphism in the HA gene, leading to the proposal that this genetic change may be influencing susceptibility and virulence in this species [24].

Our current study describes the separate inoculation of chickens and turkeys, using both *wt* and ty-ad H7N9 LPAIV isolates. The infection outcomes in both poultry hosts considered the different pathogenesis, infectivity, and transmissibility outcomes for the ty-ad and the *wt* viruses. We also explore factors which might impact on the infection outcome, including tissue tropism, receptor distribution and replication kinetics in different avian and mammalian cell types. The zoonotic and reverse-zoonotic potential of the ty-ad variant was also assessed.

## METHODS

### Viruses and the generation of a turkey-adapted (ty-ad) virus variant

Two main viruses were used in this, (i) A/Anhui/1/13 (H7N9) termed wt and (ii) a turkey adapted variant of A/Anhui/1/13 (H7N9) termed ty-ad H7N9. The H7N9 *wt* (A/Anhui/1/13 [H7N9]) was a kind gift from Professor John Macauley at the Francis Crick Institute, UK, which had undergone three egg passages (EP3) in 9-11 days-old specific pathogen free (SPF) embryonated fowls’ eggs (EFEs) [30]. Three further passages in EFEs produced the EP6 stock which served as the inocula in the *in vivo* experiments (H7N9 *wt*). The EP6 H7N9 was sequenced in its entirely and confirmed as being identical to the original GenBank sequence submission (accession numbers: CY187618-CY187625) except for an amino acid change in the HA, namely N141D (complete gene numbering). This exact virus stock was used previously to infect turkeys which led to the accumulation of amino acid substitutions [24]. A single amino acid substitution, L217Q (L226Q, H3 numbering), was consistently identified in the consensus sequence from all contact turkeys analysed [24]. To evaluate the effect of these substitutions, we previously isolated and propagated a virus (ty-ad) in 12-day-old embryonated turkeys’ eggs (ETEs) by inoculation into the allantoic cavity [24]. Whole genome sequencing (WGS) of the ty-ad isolate revealed only two polymorphisms compared to the wt H7N9 virus, both resulted in amino acid substitutions, including the L217Q substitution in the HA gene, and a T10S substitution in the neuraminidase (NA) gene.

Both viruses were titrated in EFEs to determine the 50% egg infectious dose (EID_50_) [31] and then diluted. For replication with or without trypsin, a prototypical H7N7 HPAIV was used (A/chicken/England/11406/2008 [H7N7]) as previously characterised [32]. For all viruses used to assess replication kinetics in cell culture, the virus titre was determined by plaque assay on MDCK cells as described previously using a final concentration of 1 µg/ml of TPCK trypsin (SIGMA)[22].

### Animals

For *in vivo* infection and transmission experiments, both high health status White Holland Turkeys (*Meleagris gallopavo*) and specified-pathogen-free (SPF) White-Leghorn chickens (*Gallus gallus*) were used at three-weeks of age. Birds were housed on solid flooring covered with straw, with food supplied *ad libidum* and drinking water was refreshed daily. High health status ferrets (*Mustela furo*) between 700-1000g (around three months-of-age) were used in zoonotic and reverse zoonotic experiments. All birds and ferrets were confirmed as serologically negative to H7N9 by hemagglutinin inhibition (HI) using homologous antigen on serum collected prior to infection. Oropharyngeal (Op) and cloacal (C) swab (chickens and turkeys) and nasal wash (ferrets) samples collected prior to infection were used to confirm negativity to influenza A virus M-gene RNA by reverse transcription Real-Time PCR (rRT-PCR) [23, 33].

### Turkey-to-turkey infection and transmission experimental design

Two separate groups of six, three-week old turkeys were directly infected (‘Donor’, D0) via the ocular-nasal route with 100 µl of phosphate buffered saline (PBS) containing 1×10^6^ EID_50_ of the *wt* or ty-ad H7N9 viruses. At one day-post-infection (dpi), six naïve (contact) turkeys (‘first recipient’, R1) were introduced into each group of directly infected turkeys. All turkeys were weighed immediately prior to infection and daily post infection.

### Intravenous turkey experimental design

The intravenous (IV) inoculation experiment was performed according to the World Organisation for Animal Health (WOAH) protocol for IV pathogenicity index (IVPI) determination [30], using three-week old turkeys, instead of chickens. Two separate groups of ten, three-week old turkeys were directly infected via the IV route with 100 µl of sterile saline containing 1×10^6^ EID_50_ of the *wt* or ty-ad H7N9 viruses.

### Chicken infection and transmission chains design

Two separate groups of six chickens were infected via the ocular-nasal route with 1×10^6^ EID_50_ of the *wt* or ty-ad viruses (D0), diluted accordingly from the respective viral stocks in sterile PBS, with 100 μl inoculum administered per bird. At one dpi, six naive chickens were co-housed with the directly infected chickens (R1). At four dpi, the D0 birds were removed, the bedding and drinking water were replaced, and six additional naive chickens were placed in contact with the R1 chickens (“second recipient”, R2) to establish a chain of transmission. Drinking water and feed were, respectively, replaced and replenished daily in both the chicken and turkey transmission experiments.

### Ferret model of zoonosis and reverse zoonosis

Six ferrets were infected via the intranasal route with 0.5 ml (0.25ml per nare) of PBS containing 1×10^6^ EID_50_ of the ty-ad virus. The experiment was conducted in two identical blocks. In each block three ferrets were housed in the middle of three conjoined housing unit, in a linear arrangement, with the central unit separated from its two adjacent units by means of a perforated perspex barrier. One ferret from each block (total of two ferrets) were culled at six dpi for immunohistochemistry (IHC) analysis. For each ferret the external nasal tissues were washed with 1 ml PBS on two, three, four, five, six, seven and eight dpi; 1 ml of PBS was used to wash the internal nasal cavity on days two, four, six and eight dpi as previously described [22, 34]. Blood was collected from all remaining ferrets at the end of the experiment at 14 dpi, for serological testing. Six chickens and six turkeys were housed in the two adjacent cages on three and five dpi for six hours, after which the birds were moved to a new room with a separate pen for each species and group. All birds were swabbed daily and monitored for clinical signs for 14 days from last exposure (dpe). Blood samples were collected from all birds exposed to ferrets at 14 dpe.

### Sample collection and processing during the avian infection / transmission studies

All turkeys and chickens were swabbed once daily from the Op and C cavities and swabs processed into 1 ml of brain heart infusion broth (BHIB) as previously described [24]. Tissue samples were collected from two turkeys and two chickens pre-destined (random allocation), from each D0 group, for culling while apparently healthy on four and six dpi. All tissue samples were taken into 1 ml of BHIB to achieve around 10 % (w/v) for each sample. Bloods were collected via wing bleeds from live birds during the study, and via terminal heart bleeds from surviving birds at study-end, with bleed times indicated in the results and relevant figures. Samples of drinking water and faeces were collected from the environment of infected or contact birds and processed as previously described [35].

### RNA extraction and AIV rRT-PCR

RNA was extracted from swabs, tissues and environmental samples as previously described [24] and extracted RNA was tested by the M-gene rRT-PCR which featured the primers and probe designed by Nagy, Vostinakova [33], as described previously to detect AIV viral RNA (vRNA) [24]. Ct values were titrated against a ten-fold dilution series of H7N9 *wt* RNA of a known titre and were converted to relative equivalent units (r.e.u.) as described previously [23]. Ct values ≥ 36 were considered negative for influenza A virus.

### Whole genome sequencing (WGS)

Whole-genome sequencing was undertaken on the RNA extracted from inocula, and clinical samples as previously described [24]. Briefly, double-stranded complementary DNA (cDNA) was synthesized using the cDNA synthesis system (Roche, UK) following the manufacturer’s protocols. Quantification of the synthesized cDNA was performed utilizing the fluorescent PicoGreen reagent, and 1ng of the cDNA was employed as a template for the construction of the sequencing library using the NexteraXT kit from (Illumina, Cambridge, UK). Subsequently, the sequencing libraries were processed on a MiSeq instrument (Illumina, Cambridge, UK), generating paired-end reads of 2×75 bases. Genetic analysis was performed using MEGA X (Mega Software) [36]

### Serology

Serological analysis was undertaken as previously described for sera of turkey [24] and chicken [23]. Ferret sera were treated with four volumes of receptor-destroying enzyme (APHA Scientific, Weybridge, UK) as previously described [22]. Detection of seroconversion to subtype-specific (homologous) HA antigens was conducted using the haemagglutination inhibition (HAI) assay, using four hemagglutination units of the corresponding H7N9 LPAIV as the antigen [30].

### Replication kinetics and plaque assay

All cells were incubated at 37 °C with 5 % CO_2_ unless stated. Replication kinetics were assessed in chicken, turkey, or duck embryo fibroblasts (CEF, TEF, DEF respectively), MDCK (canine), CACO2 (human) and NPTr (swine) cells. Cells (80-90% confluency) were used in six well plates; after removal of media and wash steps with PBS, the cells were infected with 400 µl of virus at an MOI of 0.01. Each plate was then placed in an incubator for one hour, gently rocking the plate every 20 minutes. The viral inoculum was then removed, and cells washed with PBS. Dulbecco’s Modified Eagle Medium (DMEM) containing TPCK trypsin (SIGMA) with a final concentration of 1 µg/ml was then added to each well. The plates were incubated, and samples collected at indicated time points and stored at −80 °C for further use. For plaque assays, 6 well plates of TEFs or MDCKs were inoculated with 10-fold serially diluted virus in DMEM and added to the plates as above. Flu overlay (immunodiffusion grade melted agarose, Sigma-Aldrich) containing TPCK trypsin with a final concentration of 1 µg/ml was added to each well, 2 ml of this mixture was then added to each well. The plates were inverted to prevent condensation build up in the wells and incubated for a period of four days. Plaques were visualised using crystal violet (Sigma), counted and plaque forming units per ml (p.f.u. ml^-1^) were calculated as previously described [22].

### Immunohistochemistry (IHC) and characterising −2,3 and −2,6 receptor distribution in avian tissues

Tissue samples were fixed in 10 % (v/v) neutral buffered formalin (VWR, East Grinstead, UK) and processed routinely through graded alcohols and chloroform and embedded in paraffin wax. Four-micrometre-thick sections, cut on a rotary microtome, were used for immunohistochemical (IHC) detection of influenza A nucleoprotein as previously described [24]. For lectin staining of virus receptor distribution in tissues, slides were dewaxed in xylene and passed through graded alcohols to Tris-buffered saline solution with 0·05% Tween (TBSt) (0·005M Tris, pH 7·6, 0·85% w/v NaCl). Samples were subsequently incubated with biotinylated *Maackia amurensis* lectin II (1/100) (Vector Laboratories, Peterborough, UK) for the α2,3 ‘avian receptor’, or biotinylated *Sambucus nigra* (Elderberry) bark lectin (1/1000) (Vector Laboratories) for the α2,6 ‘human receptor’ for 1 hour, and VECTASTAIN Elite ABC-peroxidase reagent (Vector Laboratories) for 30 min, at room temperature. Sections were washed three times with TBSt between incubations. The IHC staining was visualised using 3,3 diaminobenzidine (Sigma-Aldrich, Poole, UK), and sections were counterstained in Mayer’s haematoxylin (Surgipath, Peterborough, UK), dehydrated in absolute alcohol, cleared in xylene and mounted using dibutyl phthalate xylene (DPX) and glass coverslips.

Samples were processed and images captured in a double-blinded fashion and scored semi-quantitatively by trained veterinary pathologists based on the intensity and distribution of the staining (-, absent; +/-, minimal; +, mild; ++, moderate).

### Statistical analysis

To assess disparities among various groups, one-way analysis of variance (ANOVA), Two-way ANOVA, or paired students t-tests were carried out (GraphPad Prism v8). A significance threshold of P values (<0.05) was set. Area under the curve (AUC) analysis was performed on total shedding as previously described [35] using (GraphPad Prism v8).

## RESULTS

### The ty-ad H7N9 virus has increased pathogenicity for turkeys but not chickens

Both the *wt* and ty-ad H7N9 viruses were previously characterised as LPAIV by standard IVPI testing in chickens [24]. Following direct ocular-nasal infection of turkeys, the *wt* virus caused 33 % mortality in the D0 turkeys between 8 and 8.5 dpi (Fig. 1A). Comparatively, the ty-ad virus caused 100 % mortality in D0 turkeys between 6 and 8 dpi. Similarly, following transmission to naive contact turkeys, the *wt* virus caused 0 % mortality, yet the ty-ad virus caused 100 % mortality (Fig. 1A). Were mortality was observed, turkeys typically exhibited mild clinical signs including ruffled feathers and huddling behaviour during the preceding timepoints. Importantly, all surviving *wt* virus infected turkeys demonstrated seroconversion to H7N9 when tested by HAI at 14 dpi which confirmed infection (data not shown). A similar difference in pathogenicity was observed when turkeys were inoculated with either virus intravenously; the *wt* virus caused 60 % mortality by 8 dpi, whereas the ty-ad virus caused 100 % mortality within 5 dpi (Fig. 1B). The ty-ad virus infected turkeys also exhibited reduced, but not statistically significant weight gain compared to the *wt* infected turkeys, for both D0 and R1 groups (Fig. S1). By contrast, chickens (both D0 and R1 groups) exhibited 0 % mortality (Fig. 1C) with either virus. Taken together, these data indicate that the ty-ad virus is more pathogenic than the *wt* virus in turkeys, but not in chickens.

**Fig. 1.**
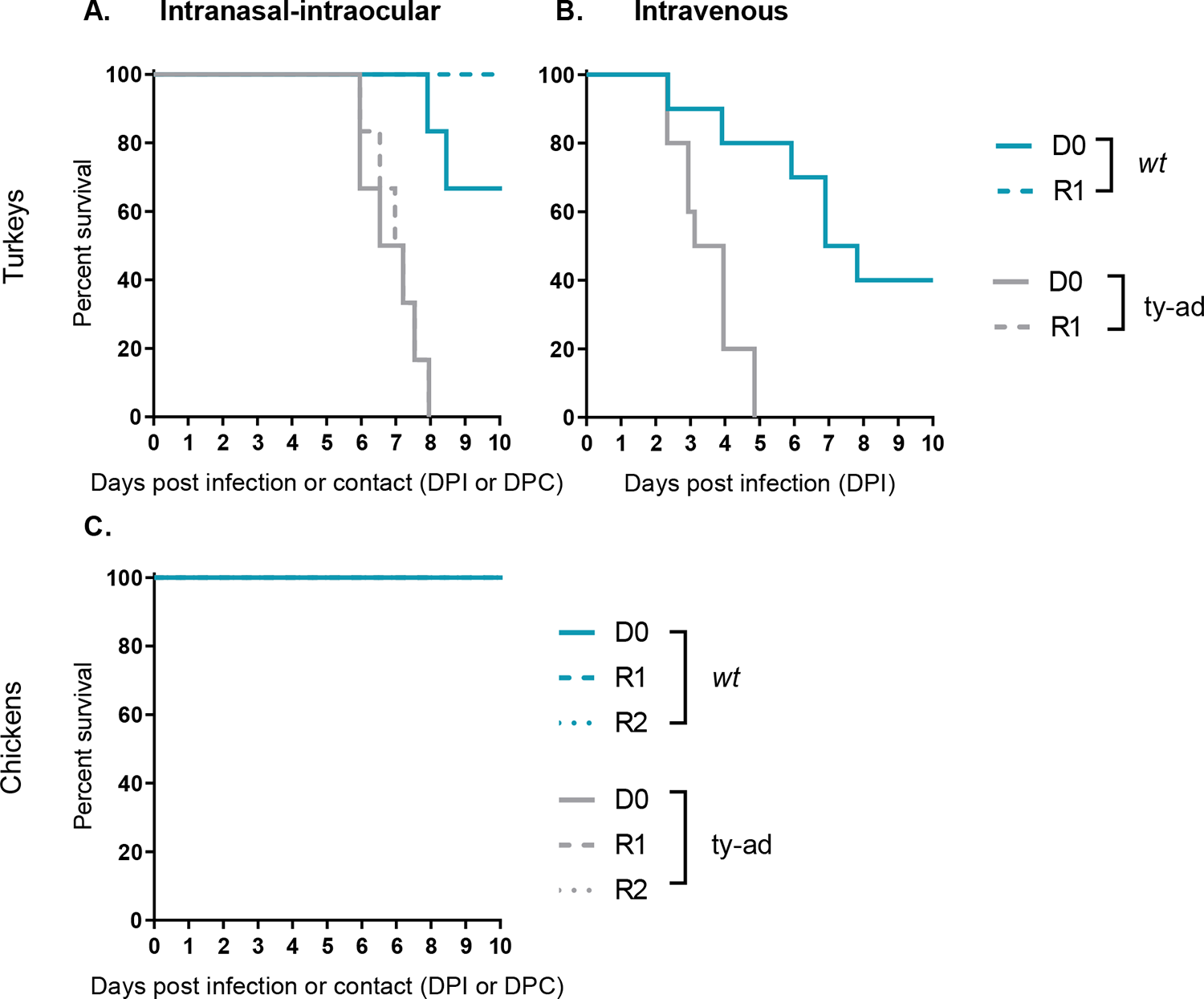
Survival of turkeys and chickens infected with H7N9 *wt* and ty-ad virus variants. **A and C.** The survival rates of six turkeys (**A**) or chickens (**C**) directly inoculated (D0) via the intranasal-intraocular route (solid lines) with 1×10^6^ EID_50_ of the *wt* (teal lines) or ty-ad (grey lines) virus variants. The survival of six turkeys (**A**) or chickens (**C**) co-housed with either group at 1 day post infection (R1) (dashed lines) or six contact chickens added at 4 dpi (dotted lines) is also shown. **B.** The survival rates of 10 turkeys inoculated intravenously with 1×10^6^ EID_50_ of *wt* (teal lines) or ty-ad (grey lines) virus variants.

### Both the *wt* and ty-ad virus infected birds shed similar levels of vRNA and transmitted infection equivalently in both turkeys and chickens

Both the *wt* and ty-ad viruses successfully infected all six of the respective D0 turkeys or chickens. For both groups, vRNA was detected in both the Op and C swabs taken from turkeys (Fig. 2A and B) and chickens (Fig. 3). Directly infected (D0) turkeys and chickens shed peak levels of vRNA of between 1×10^5^ to 1×10^6^ r.e.u. from the Op cavity following infection with either virus (Fig. 2A and Fig. 3A and C). Based on AUC analysis, there was no significant difference in the total amount of virus shed from turkeys following infection (Fig. 2E and F). The levels of vRNA detected in the water, and environmental faecal sources, were largely comparable between turkeys and chickens infected with either virus, although greater vRNA was detected in the faeces deposited by the ty-ad group between 4 and 8 dpi (Fig. S2B).

**Fig. 2.**
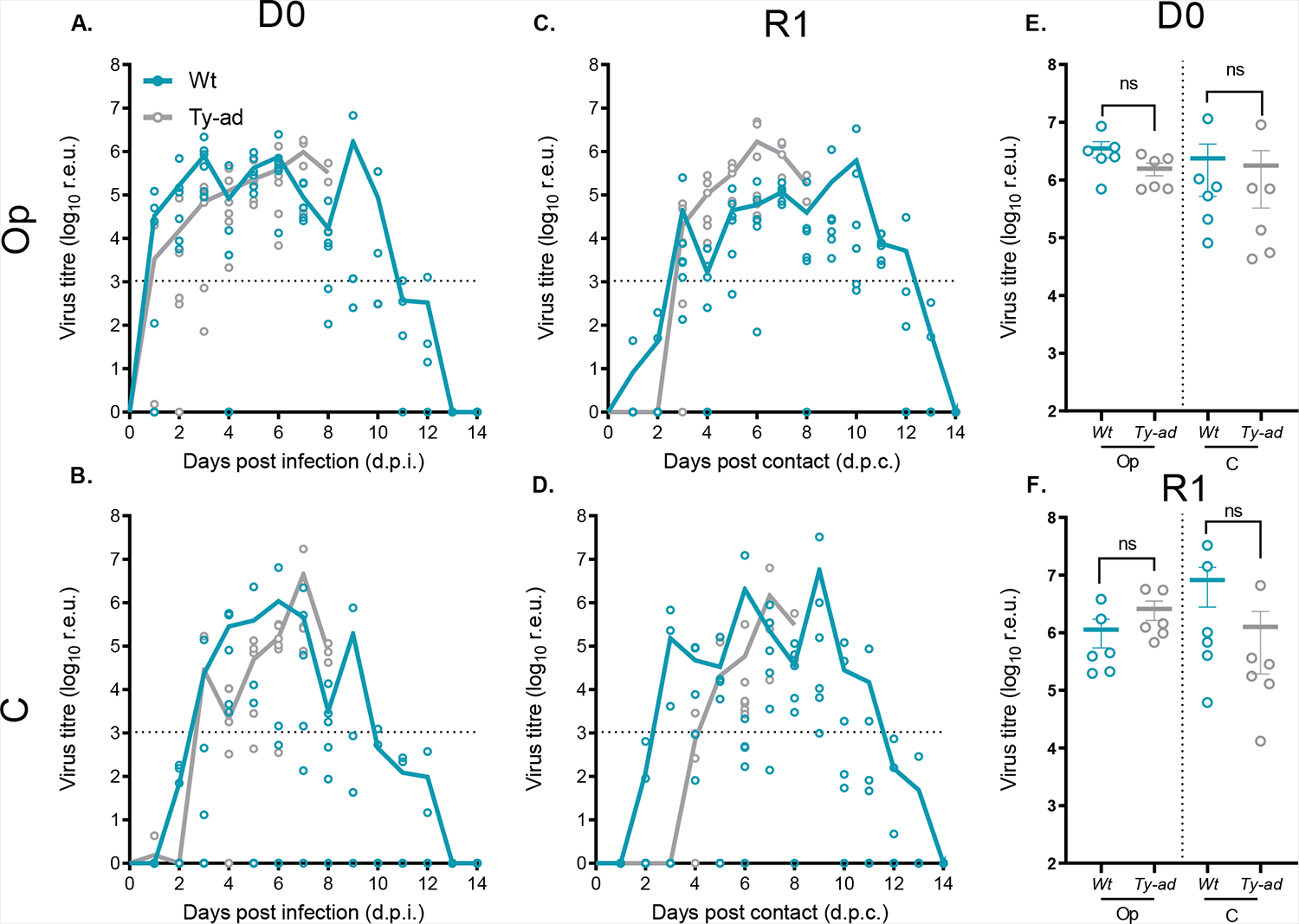
Viral RNA shedding from *wt* and ty-ad virus infected or contact turkeys. Viral RNA shedding profiles from turkeys directly inoculated with 1×10^6^ EID_50_ with *wt* (teal lines) or ty-ad (grey lines) virus variants (D0) (**A & B**) or turkeys co-housed with either group at 1 day post infection (R1) (**C & D**). Symbols show individual bird vRNA levels and lines indicate the mean vRNA levels for all live birds in a given group, quantified as r.e.u. (see Methods for details). Viral RNA shedding was quantified from swabs collected from the Op (**A & C**) or C cavities (**B & D**). Dotted horizontal lines indicate the limit for positivity at 36 ct values (1.05×10^3^ r.e.u). Shedding profiles from individual birds were used to calculate area under the curve (AUC) and graphically displayed for the directly infected (D0) (**E**) or contact (R1) (**F**) birds. One-way ANOVA with multiple analysis was performed comparing the Op and C AUC shedding for both the D0 and R1 groups, ns (not significant) indicates P-value >0.05.

**Fig. 3.**
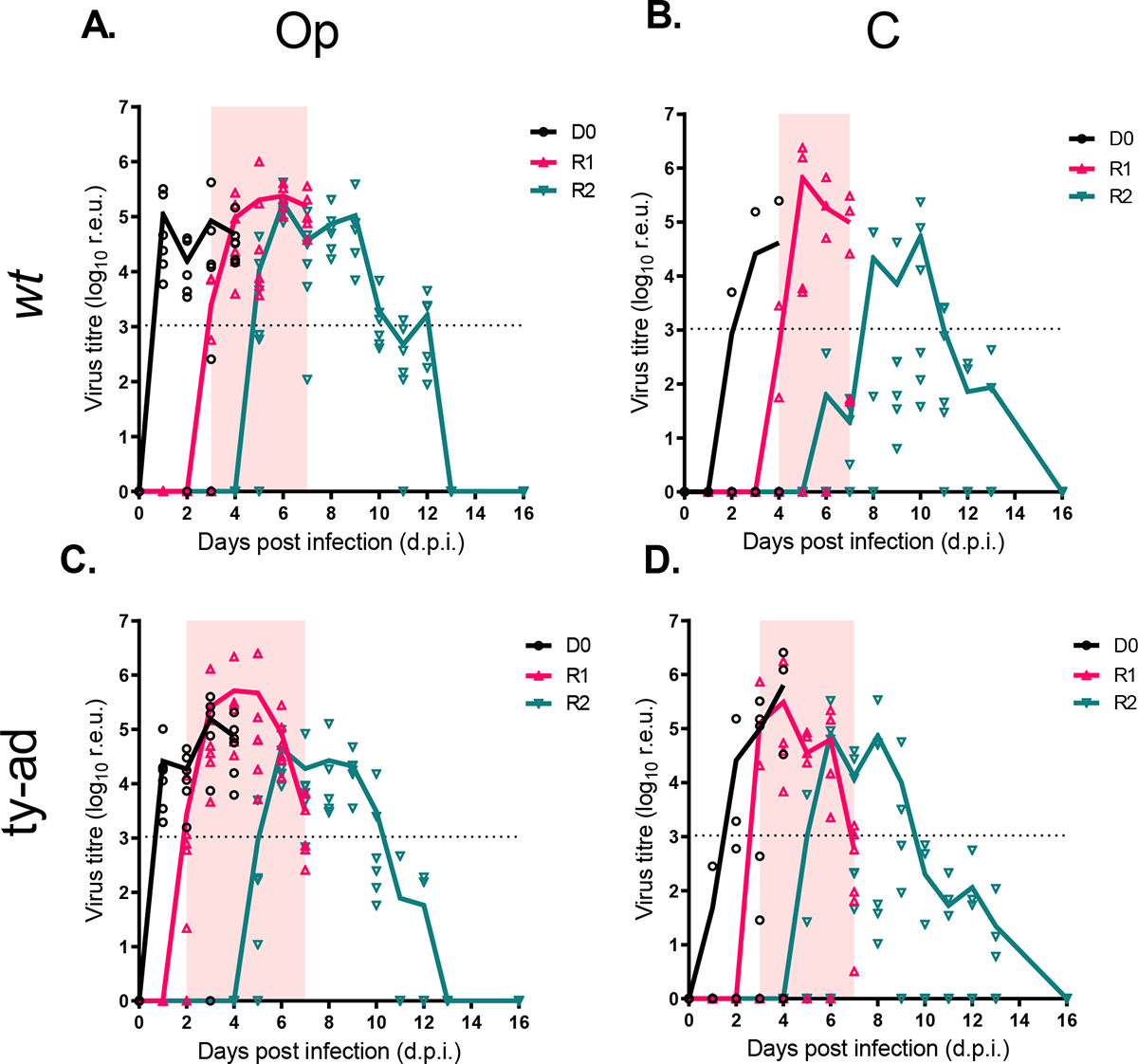
Viral RNA shedding from *wt* and ty-ad virus infected or contact chickens. Viral RNA shedding profiles from chickens directly infected with 1×10^6^ EID_50_ with *wt* (**A & B**) or ty-ad (**C & D**) virus variants (D0) (black lines) or chickens co-housed with either group at 1 dpi (R1, red lines) or 4 dpi (R2, teal lines). The D0 chickens were removed prior to the addition of second contact (R2) chickens at 4 dpi to facilitate an extended transmission chain. Symbols relate to individual birds and lines indicate the mean virus titre (r.e.u.), the pink shaded areas indicate the window of detectable viral RNA being shed from the R1 chickens. Dotted horizontal lines indicate the limit for positivity at 36 ct values (1.05×10^3^ r.e.u). Viral RNA shedding was quantified in from swabs collected from the Op (**A & C**) or C cavities (**B & D**).

The transmission efficiency was 100 % for both viruses in both species; all R1 turkeys and chickens became infected and shed vRNA from the Op and C cavities (Fig. 2C & D and Fig. 3). For the turkeys, positive vRNA levels were first detected for both *wt* and ty-ad R1 groups at 3 days post-contact (dpc), indicating transmission within a similar timeframe. Op and C vRNA shedding profiles for the R1 contact turkeys were similar to those of the directly infected turkeys (Fig. 2A and B compared to C and D). For the chickens, a chain of transmission was established from D0 to R1 and subsequently to R2 chickens, with 100% successful transmission throughout the chain. However, the ty-ad vRNA was detected in swab material from the R1 chickens a day earlier than the *wt* virus (Fig. 3A and C).

All chickens and turkeys which survived until the end of the study (19 dpi) were seropositive by the HAI against the H7N9 antigen (data not shown). Interestingly, using post infection sera raised against the *wt* or ty-ad viruses, there was a statistically significant drop in reciprocal HAI titres between the wt (L217) and ty-ad (Q217) antigens, for heterologous compared to homologous sera, suggesting that aa position 217 may be an antigenically important residue (Fig. S3).

### Differences in systemic H7N9 distribution between turkeys and chickens, with increased tissue dissemination of ty-ad in turkeys compared to *wt* virus

The tropism of the *wt* and ty-ad viruses was investigated by vRNA quantification and viral antigen detection by IHC. Tropism was investigated in tissues collected from at least two turkeys and two chickens per group - culled while apparently healthy on 4 and 6 dpi (Fig. 4). In the turkeys, both the *wt* and ty-ad vRNA were detected in a range of organs, including respiratory (nasal turbinates, trachea lung and air sacs), and other visceral (including the intestine, pancreas spleen, heart, liver and brain) organs (Fig. 4A and B, Fig. S4).

**Fig. 4.**
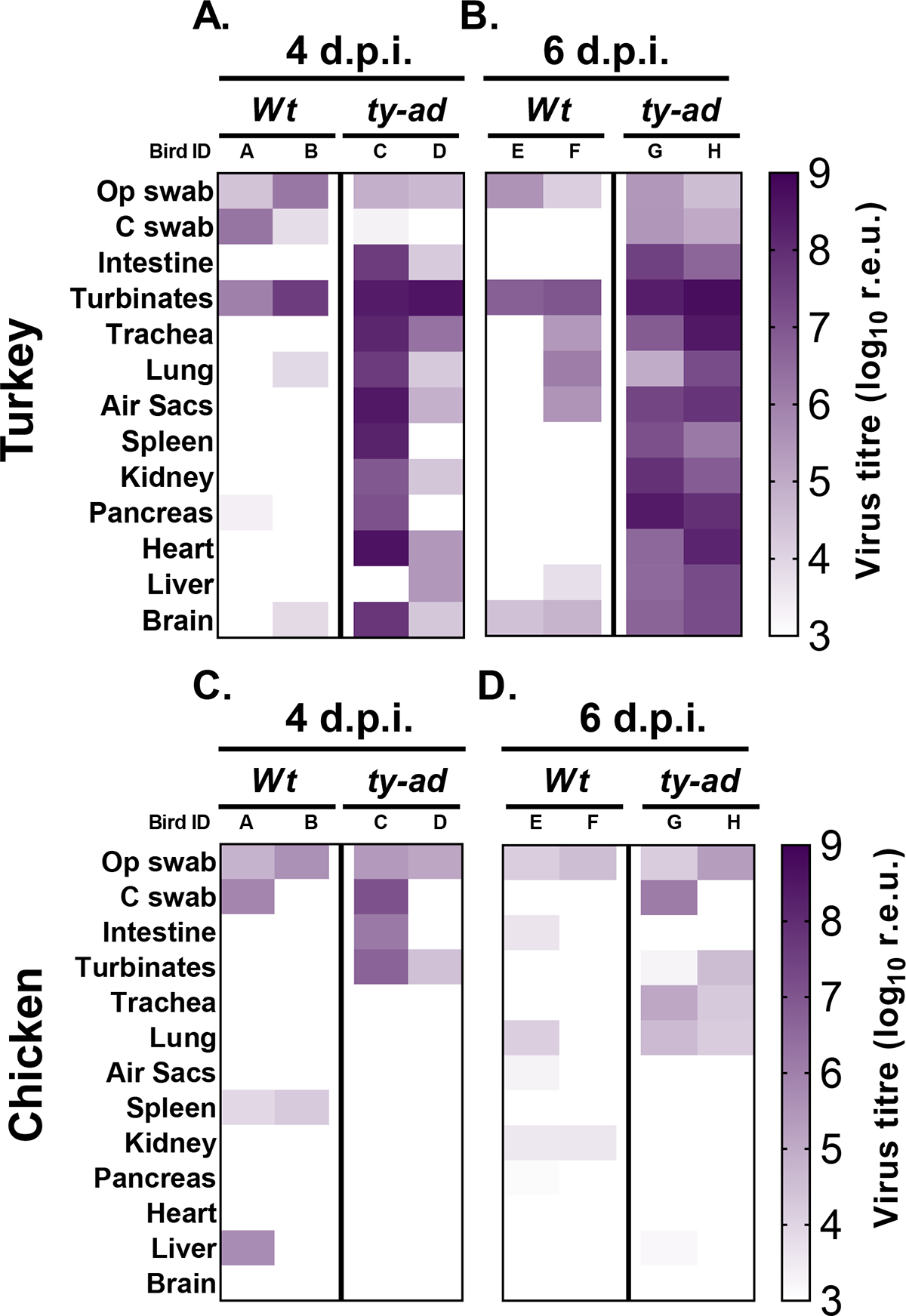
Viral RNA levels in different organs of directly infected turkeys and chickens. Viral RNA was quantified in organs collected from turkeys (**A & B**) or chickens (**C & D**) directly infected with 1×10^6^ EID_50_ of the *wt* or ty-ad virus variants. Organs were taken from 2 birds in each group which were culled at 4 dpi (**A & C**) or 6 dpi (**B & D**). Detection of viral RNA is shown graphically in a heat map ranging from the limit of positive detection at 1.05×10^3^ (pale colour) to 1×10^9^ r.e.u. (dark colour).

However, the ty-ad virus was detected more frequently and with higher vRNA and IHC (Table S1) levels than the *wt* virus. By contrast, chickens exhibited considerably lower vRNA levels in all organs, with limited dissemination beyond the respiratory and enteric tracts, at both 4 and 6 dpi (Fig. 4A and B compared to D and E). There were also no major differences in virus tropism between the *wt* and ty-ad virus in the chickens (Fig. 4C and D, Table S1).

### Turkeys and chickens have slightly different sialic acid receptor distributions in their respiratory tracts and visceral organs

The L217Q mutation has been previously shown to change virus receptor preference from α-2,6-sialyllactosamine (α-2,6) to α-2,3-sialyllactosamine (α2,3) [12]. The ty-ad virus possesses a Q at amino acid position 217, therefore we hypothesised that differences in α-2,3 and α-2,6 linked SA might explain the expanded tissue tropism of the ty-ad virus in turkeys. Using lectins which bind to α-2,3 (Maackia Amurensis Lectin II [MALII]) or α-2,6 (sambucus nigra [SNA]), the host receptor distribution was assessed in turkey and chicken tissues where the highest levels of vRNA were detected (nasal turbinates, trachea, lung, pancreas and kidney) (Fig. 4 and S5, Table 1). Turkey and chicken tissues had similar α-2,3 and α-2,6 receptor staining distribution, although some differences were observed. Turkey tissues had more intense MALII (α2,3) labelling in the nasal cavity and trachea compared to SNA (α2,6) staining (Table 1). Further, in turkeys staining in the nasal cavity and trachea was more intense for MALII (α-2,3) compared to that observed in the same tissues from chickens (Table 1). Interestingly, whilst SNA (α-2,6) staining in the trachea of turkeys was less pronounced than that seen in chickens, the SNA (α-2,6) staining in turkey kidneys was more intense than observed in chickens (Table 1).

**Table 1.** α2,3 and α2,6 linked sialic acid distribution in chicken and turkey tissues.

| Lectin (receptor preference) | Tissue | Chicken |  |  | Turkey |  |  | Chicken consensus | Turkey consensus |
| --- | --- | --- | --- | --- | --- | --- | --- | --- | --- |
|  |  | A | B | C | A | B | C |  |  |
| MAL II (α2,3) | Nasal cavity | +/- | + | + | ++ | ++ | ++ | + | ++ |
|  | Trachea | +/- | + | + | ++ | ++ | ++ | + | ++ |
|  | Lung | + | + | + | + | + | + | + | + |
|  | Pancreas | + | + | + | + | + | + | + | + |
|  | Kidney | ++ | ++ | ++ | ++ | ++ | ++ | ++ | ++ |
| SNA (α2,6) | Nasal cavity | + | ++ | + | +/- | + | + | + | + |
|  | Trachea | + | ++ | ++ | + | ++ | + | ++ | + |
|  | Lung | + | + | + | + | + | + | + | + |
|  | Pancreas | +/- | + | + | ++ | + | + | + | + |
|  | Kidney | + | + | + | ++ | ++ | ++ | + | ++ |

Semi-quantitative scoring was performed in blinded fashion; -, absent; +/-, minimal; +, mild; ++, moderate

### Both *wt* and ty-ad viruses require trypsin for efficient replication, but the ty-ad virus replicates to higher titres in avian cells

vRNA from Op swab samples collected at six dpi from all six turkeys directly infected with the ty-ad virus was sequenced. Sequence analysis revealed a single basic cleavage site (CS) sequence present in HA (PEIPKGR’GLF) in all sequences, identical to the CS sequence in the wt and ty-ad viruses used for the inocula. In cell culture, both the *wt* and ty-ad H7N9 viruses required the presence of exogenous TPCK trypsin to replicate; both viruses formed plaques when TPCK trypsin was added yet failed to form plaques without TPCK trypsin addition (Fig. 5A). In comparison, a H7N7 HPAIV formed plaques even without trypsin (Fig. 5A). The *wt* and ty-ad viruses were also compared for replication in primary duck, chicken and turkey embryonic fibroblast (DEF, CEF and TEF) cells as well as human (CACO-2), swine (NPTr) and canine (MDCK) cells (a cell line highly susceptible and permissive to IAV replication). In all three of the different avian cells, the ty-ad virus replicated to statistically significantly higher titres compared to the *wt* virus (Fig. 5B-D). In the different mammalian cells however, there was no statistically significant difference in virus replication between the *wt* and ty-ad viruses over the 72-hour culture period (Fig. 5E-G).

**Fig. 5.**
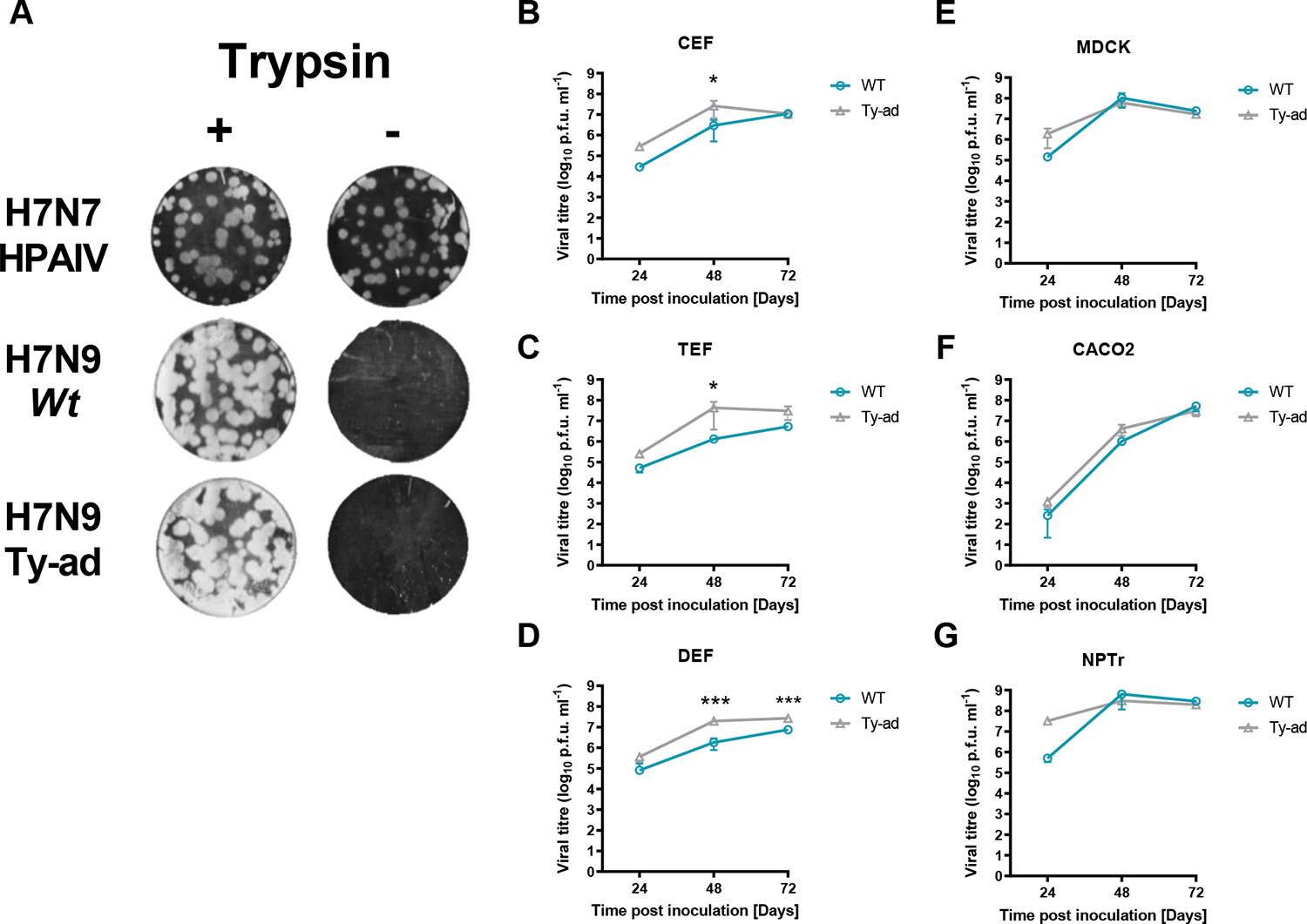
Replication kinetics of *wt* and ty-ad virus variants in cells from different species. **A.** Representative well visualising plaques on turkey embryo fibroblasts (TEF) cells from three independent replicates with the same dilution of wt, ty-ad or A/chicken/England/11406/2008 (H7N7 HPAIV) viruses in the presence (+) (1µg/ml) or absence (-) of TPCK trypsin. Plaque assays were stained with crystal violet. **B-G.** Virus replication of *wt* (teal) and the ty-ad (grey) virus variants in chicken, turkey or duck embryo fibroblasts (CEF, TEF, DEF), MDCK (canine), CACO2 (human), NPTr (swine) cells. Cells were infected at a MOI of 0.01 with each virus and cell supernatant was harvested at 24, 48 or 72 hours post infection. Viral titres were determined based by plaque forming units per ml (p.f.u. ml^-1^) using plaque assay. Each time point corresponds to the mean of four biological replicates with standard deviations indicated. Two-way ANOVA with multiple analysis was performed comparing the two strains, * indicates P-value <0.05, ** indicates P-value < 0.005, *** indicates P-value ≤ 0.001.

### The ty-ad virus represents a zoonotic threat but does not exhibit reverse zoonotic transmission under experimental conditions

We next investigated the zoonotic consequence of the ty-ad virus by investigating its ability to infect and replicate in ferrets, a model for human influenza A virus infection. Four ferrets (two per block) were directly inoculated intranasally with 1×10^6^ EID_50_ of the ty-ad virus and successfully became infected, with vRNA being detected in interior nasal wash samples from 2 until 6 dpi (Fig. 6A). Viral RNA was also detected (above the threshold for assay positivity) from 2 to 6 dpi from exterior nasal wash samples. WGS was performed on the viruses present in the interior nasal wash samples from all ferrets at 4 or 6 dpi (Fig. 6; yellow filled circles). Sequence analysis revealed that viruses present in all ferrets showed no polymorphisms compared to the input virus at a consensus level (data not shown). Influenza viral NP was also detected in the respiratory turbinates, trachea and lungs of the two ferrets pre-destined for IHC analysis at six dpi (Fig. S6 and Table S2). All four of the remaining ferrets bled at study-end (14 dpi) had seroconverted to homologous H7N9 antigen by HAI (Fig. 6B). To mimic reverse zoonoses, the ability of infected ferrets to transmit virus to naive turkeys and chickens was investigated. No vRNA was detected in either avian Op or C swabs collected daily for 14 dpe from either block (data not shown). In addition, at 14 dpe, none of the chickens or turkeys had seroconverted to homologous H7N9 antigen by HAI from either block (Fig. 6B).

**Fig. 6.**
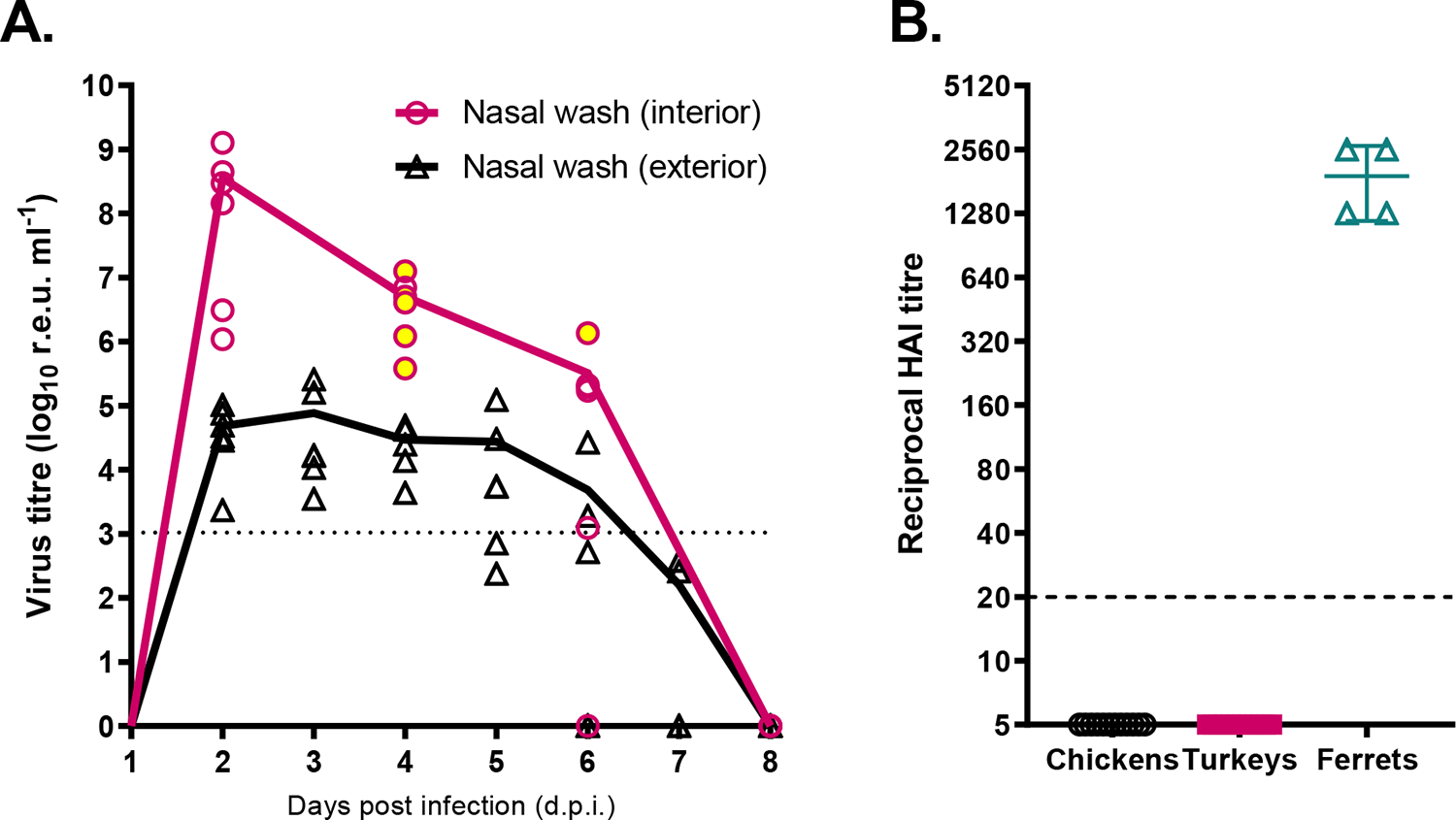
Zoonotic and reverse zoonotic infection dynamics for the ty-ad virus. Infection dynamics of the ty-ad virus in the ferret model and investigation of reserve zoonotic infection from ferrets to chickens and turkeys. **A.** Virus titre estimates (r.e.u.) derived from interior (pink) or exterior (black) nasal wash samples collected from six ferrets (three per block) directly infected intranasally with 1×10^6^ EID_50_ of the ty-ad virus. One ferret per block (total of two ferrets) were culled at 6 dpi for IHC analysis. Symbols show an individual ferret’s vRNA levels in the washes, while continuous lines indicate the mean vRNA levels. Yellow filled circles indicate the interior nasal wash samples for which sequencing of amino acid position 217 in HA was performed and confirmed as Q217 (samples from five ferrets at four dpi and one ferret at six dpi). Dotted horizontal line indicates the limit for positivity at 36 ct values (1.05×10^3^ r.e.u) **B.** HAIs were performed on sera taken from four directly inoculated ferrets at the end of the experiment (14 dpi). Chickens (n=12) and turkeys (n=12) were similarly bled at 14 dpi, after having been housed in the same air space, yet physically separated, from the infected ferrets for a 6 h period at 3 dpi. HAI assays were performed using homologous H7N9 antigen. Dashed horizontal line indicates the limit for positivity at a titre of 20 HAI values (1.05×10^3^ r.e.u).

## DISCUSSION

Since its emergence in 2013 in China, A/Anhui/1/13-lineage H7N9 LPAIV has continued to circulate in poultry threatening human health. Currently H7N9 is located exclusively in China [37] where poultry production consists largely of chickens and ducks with almost negligible turkey production [25]. Despite widespread poultry vaccination in China, H7N9 continues to be detected in poultry [3], with an ever present concern that geographical expansion of H7N9 could increase its zoonotic and epizootic risk through reassortment with indigenous AIVs, or access to novel hosts [22].

Previously, the risk of A/Anhui/1/13-lineage H7N9 infection in turkeys was investigated, modelling a scenario in which H7N9 virus expands into neighbouring countries with a larger turkey poultry sector [24]. Infection and transmission of H7N9 LPAIV in turkeys was associated with the 217 L217Q (H7 numbering of the mature protein) polymorphism in the HA gene [24]. An H7N9 virus from a turkey with severe clinical disease (termed the ‘ty-ad’ virus) was isolated that contained two amino acid substitutions, one in the HA (L217Q) and one in the NA (T10I) proteins. In this study we explored the outcome of infection for two gallinaceous poultry species (chicken and turkey) with this ‘ty-ad’ virus variant compared to the *wt* virus. We further investigated potential mechanisms which might underpin the differences in observed infection outcome and explored the zoonotic and reverse zoonotic risk of these viruses.

*In vivo,* when inoculated intravenously or intranasally, the ty-ad virus caused greater, and more rapid, mortality in turkeys than the *wt* H7N9 virus strain. This disparity in clinical disease between the *wt* and ty-ad virus was not observed in experimentally infected chickens. Post-mortem investigation of the turkeys in the ‘ty-ad’ virus infected group revealed a systemic tissue dissemination of vRNA with higher levels of viral RNA being detected in a wider range of visceral organs compared to the *wt* virus. By contrast, following chicken infection, this difference in vRNA levels and its systemic distribution was not observed between the two virus variants. However, both viruses displayed comparable shedding profiles from both the Op and C cavities. Different pathogenic outcomes between chickens and turkeys infected with H7 AIVs has been reported previously [38]. However, the genetic determinants for virulence and pathogenesis of HPAIV in turkeys are largely unknown. For AIVs, tropism is determined by a combination of factors including (i) the distribution of proteases required to cleave HA0 to HA1 and HA2 required for infectivity, (ii) the distribution of the sialic acid (SA) receptors to which the viral hemagglutinin (HA) binds, and (iii) cells being permissive (allowing viral replication) for viral infection. We therefore explored these factors which might underpin the observed differences in pathogenicity and tropism, between the turkey and chicken hosts following infection with the two H7N9 LPAIV variants.

The major genetic determinant of AIV tropism is the presence of a multiple basic amino acids proximal to the cleavage site in the HA protein, termed a multibasic cleavage site (MBCS). This MBCS facilitates the cleavage of HA0 to HA1 and HA2 (required for infectivity) by proteases found ubiquitously in the animal tissues [39]. Sequence analysis of the *wt* and ty-ad virus inocula confirmed a single basic cleavage site typical of a LPAIV. Moreover, both viruses required TPCK trypsin for growth in turkey cells (TEFs), affirming a LPAIV phenotype. Therefore, the expanded tropism of the ty-ad virus in turkey tissues may be independent of the restriction imposed by the tissue localisation of proteases, which typically restricts the tropism of most LPAIVs to mainly the respiratory and / or enteric tracts of chickens and turkeys [39].

Proteolytic activation of LPAIV HA occurs extracellularly driven by airway proteases, the trypsin-like serine proteases human airway trypsin-like protease (HAT) and transmembrane protease serine 2 (TMPRSS2), albeit a minority of viruses possess HA cleavage site motifs that are processed by other proteases (reviewed by Kido, Okumura [39]). These ‘atypical’ viruses, including the 1918 influenza A virus [40], can utilise proteases found ubiquitously in tissues of multiple organs for cleavage, and therefore can potentially increase tissue dissemination and pathogenicity. For H9N2 infections, changes in the cleavage site have been shown to alter the clinical outcome in turkeys [41]. The predominant proteases present in other cell types and how they may differ in distribution in turkeys compared to chickens, along with which proteases may differentially cleave the HA of the *wt* and ty-ad viruses, remain to be identified. However, conserved expression and functionality of the furin protease has been observed between chickens and ducks [42].

Therefore, considering that turkeys and chickens are closely related galliformes, with ∼90% genome identity [43], it is unlikely that notably separate protease compositions exist between the two species. However, non-basic amino acid variations in H9N2 viruses have been shown to bestow a HPAIV phenotype in turkeys compared to chickens [41]. However, the L217Q substitution, present in the HA of the ty-ad virus, is not located near to HA1/HA2 cleavage site and is therefore unlikely to alter the protease preference.

The L217Q polymorphism has been previously shown, for various IAV strains including H7N9, to change the receptor preference by increasing the affinity of haemagglutinin to α-2,3 SA [12–14]. The A/Anhui/1/13 isolate being a human derived virus has a naturally high affinity to α2,6, but is also able to bind α-2,3 SA with a relatively high affinity [14, 44–46]. Recent receptor preference analyses for H7N9 have demonstrated that Q217 increases affinity to α2,3 SA [14]. Consequently, we investigated whether the expanded tropism could be due to a greater density of α2,3 SA in turkey tissues. In this study we did not see any difference in pathogenicity between the wt and ty-ad viruses in chickens, we therefore compared the receptor distributions between turkey and chicken tissues. Using lectins which bind preferentially to α2,3 or α2,6 SA, we found that turkeys have a greater abundance of α2,3 SA in the nasal turbinates and trachea, comparatively higher than those found in equivalent chicken tissues. The nasal turbinates and trachea are the major site of H7N9 replication, and therefore might explain the rapid emergence of the L217Q polymorphism reported previously [24]. However, analysis of the turkey visceral organs revealed comparatively lower α2,3 levels than α2,6 SA, as observed by others [47], suggesting that differences in receptor distribution alone are not likely responsible for the increased tropism or observed pathogenicity.

Receptor distribution in chicken and turkey tissues performed by lectin binding served to assess the overall distribution of the α2,3 and α2,6 sialosides, and did not identify other receptor derivatives, such as those possessing specific modifications, e.g. sulphated α2,3 sialosides which appear to be important as receptors for other AIVs [48, 49]. It has also been recently reported that the abundance of N-linked glycans on the cell surface may also influence the receptor-binding preference of AIVs [50]. Therefore, a more detailed investigation into the glycome of different avian species is required to fully explore this mechanism, and its possible role in effecting H7N9 LPAIV binding to different turkey tissues, and how this may differ for the H7N9 wt and ty-ad LPAIV variants.

We also compared the replication kinetics of both the *wt* and ty-ad viruses in a range of avian and mammalian cells. In turkey, chicken and duck primary cells the ty-ad virus showed a statistically significant increase in the level of replication, compared to the *wt*. However, no statistically significant differences in replication were observed in either human, swine, or the highly sensitive and permissive MDCK (canine) cell line. Together, these observations indicated that the ty-ad virus confers a generic replicative advantage in avian, but not mammalian cells. This similar level of replication in mammalian cells with Q217 has also been previously observed [14]. Clearly, multiple factors determine the replication kinetics of AIVs. Considering that the polymorphisms in the ty-ad virus are outside the polymerase, the dynamism of vRNA replication is unlikely to be a contributory factor. This observation is more likely due to different receptor utilisation, or possibly related to a different pH of HA activation. The optimal pH of HA activation, or pH of fusion, is species dependent, and pH has been demonstrated to alter replication kinetics in *ex vivo* organ cultures [51] and for an H5N1 HPAIV isolate *in vivo* [52]. The Q217 slightly decreases the pH of fusion when assessed by syncytia formation assays [14]. Interestingly, it has been suggested that AIVs tend to favour a lower pH of fusion upon infection and circulation in turkeys, in comparison to those isolated from other avian hosts [53].

Another consideration, not explored in this study, is the differential host response induced by the *wt* and ty-ad viruses. Indeed, turkeys appear to mount a different host response than chickens in response to H7 infection, particularly from genes involved in RNA metabolism and the innate immune response [38].

However, it is uncertain whether single amino acid changes in the HA and NA proteins would contribute directly to different immune responses or enhance immune evasion of these viruses. The immune responses to influenza A virus infection in different avian species, including different galliforme species, is understudied and requires further investigation.

Alongside the Q217 polymorphism identified in the ty-ad virus, a threonine to isoleucine polymorphism was also detected in the NA protein at residue position 10 (T10I). The function of this mutation has not been documented in the literature. However, this polymorphism is located in the transmembrane domain of NA, away from the functional site and stalk, hence this polymorphism is unlikely to contribute to the different pathogenicity observed across the two species.

Antigenic comparisons between viruses possessing either Q217 or L217 induced 23- and 8-fold reductions in HAI titre with ferret and chicken antisera, respectively [54]. In this study we saw similar antigenic differences between the ty-ad (Q217) and *wt* (L217) H7N9 viruses, reinforcing this finding. However, the differential receptor binding of the L217Q mutation has been postulated to reduce the avidity of binding to chicken red blood cells, thereby over exaggerating antigenic changes in HAI assays [12].

Despite the possible effect of different avidities, the L217Q polymorphism has previously been shown to emerge as an antigenic ‘escape’ mutant upon incubation with H7N9 A/Anhui/1/13 antisera, indicating a genuine role of Q217 in the HA antigenicity of H7N9 [54]. In addition, since 2017, China has implemented a mass vaccination program of poultry against H7N9. This campaign has successfully reduced zoonotic cases yet has not eradicated H7N9 from poultry in China. Genetic analysis of H7N9 viruses detected in poultry after the introduction of the vaccine has highlighted an increased detection of Q217 in H7N9 HA gene sequences [10, 11]. The vaccine consists of the HA antigen derived from an A/Anhui/1/13-like candidate vaccine virus [54]. As such, this work suggested that Q217 emergence through antigenic evolution may afford greater pathogenic outcomes in some susceptible avian species, such as turkeys. Through *in vitro* characterisation, Q217 appears to have reduced zoonotic ability [14]. However, in this study we observed that the ty-ad virus was still capable of infecting ferrets, producing similar levels of vRNA in nasal wash samples, when compared to that of H7N9 *wt* virus ferret infections conducted in the same animal facilities [22, 34]. This observation suggested that while the ty-ad virus possess no greater zoonotic risk to *wt*, it still likely retains its ability to infect humans. We also explored the potential for this virus to spread from infected ferrets to chickens or turkeys, in an attempt to mimic reverse zoonotic transmission. In this study, infected ferrets were physically separated from chickens and turkeys but shared the same airspace for two 6-hour periods. No evidence of airborne transmission to naïve birds could be detected. While longer or more intermate contact may have facilitated successful transmission, these results suggest that the ty-ad H7N9 does not exhibit a strong ability for reverse zoonotic transmission. However, infection spread among avians included extension of the successful chicken transmission chain to a second contact group (R2) of chickens. Daily refreshing of the drinking water and changing of the bedding at 4 dpi suggested that spread from R1 to R2 chickens did occur, with the detected viral environmental contamination by both wt and ty-ad viruses potentially contributing to the acquisition of infection, as described elsewhere [56].

In conclusion, this study has demonstrated that the ty-ad virus emerged rapidly following infection of turkeys, possibly due to high levels of α2,3 SA in the major sites of virus replication in turkeys. When this adaptation occurred, it enabled the virus to replicate systemically in a wider range of organs, and at implied higher titres. This enhanced tropism conferred increased pathogenesis and mortality to infected turkeys. The enhanced tissue tropism was not due to any change in the ability of the virus to replicate without trypsin, or due to any obviously different receptor distributions in the turkey organs. However, the adaptation did confer a statistically significant increase in replication in avian cells, including turkey cells, but not mammalian cells. The exact mechanism for the systemic tropism of ty-ad virus and its associated pathogenesis in turkeys remains elusive. However, this work highlighted the requirement for continued surveillance of low pathogenicity H7N9, particularly as vaccination programmes continue.

## Acknowledgements

The authors would like to thank Alex Byrne, Carlo Bianco and Samantha Watson for assistance with the experimental work.

## Author contribution

M.J.S., A.H.S., S.M.B. and J.J. conceived and designed the studies. J.J., A.H.S., S.M., A.N., S.T., M.J.S. performed the experimental work. J.J., S.M., S.T., M.J.S., A.C.B interpreted the data. J.J. and A.C.B, drafted the manuscript. All authors reviewed and edited the manuscript.

## Ethical and safety statement

All animal experiments and procedures required approval from the local APHA Animal Welfare and Ethical Review Body to comply with the relevant European and UK legislation and were carried-out in accord with the Home Office (UK) Project Licences 70/8332 and 70/8692. Any infected poultry which began to display severe clinical signs as described previously [55] and were euthanised and were recorded as a mortality. UK regulations categorise the H7N9 LPAIV as a SAPO-4 and ACDP-3 pathogen because it is a notifiable animal disease agent and presents a zoonotic risk. All laboratory and containment work with H7N9 specimens, including infected poultry, was done in licenced BSL3 facilities.

## SUPPLEMENTARY FIGURE LEGENDS

**Fig. S1.** Percent weight change of turkeys infected (D0) with *wt* or ty-ad virus variants, or of turkeys housed in contact (R1) with either group. Six turkeys (D0) were infected via the oronasal route with 1×10^6^ EID_50_ or either *wt* or ty-ad virus variants. Six further turkeys (R1) were co-housed with either group at 1 day post infection. All turkeys were weighted daily, and percentage weight change was calculated from their original weight prior to infection. Dotted horizontal line indicates zero percent weight change.

**Fig. S2.** Detection of vRNA in environmental drinking water and faecal samples following infection and transmission of *wt* or ty-ad virus variants in turkeys and chickens. vRNA detection in drinking water (**A and C**) or faecal samples (**B and D**) collected from the environment of infected or contact turkeys (**A and B**) or chickens (**C and D**), from the experiments shown in Fig 2 or Fig 3 respectively.

**Fig. S3.** Homologous H7 HAI titres from infected turkeys. HAI assays were performed on paired sera taken from chickens and turkeys, following initial infection via the ocular-nasal route with (**A**) *wt* (217L) (total n=17; n=8 chickens; n=9 turkeys) or (**B**) ty-ad (217Q) (total n=14; n=6 chickens; n=8 turkeys) viruses. Sera were collected between 8- and 19-days post infection. The HAI assays were performed using homologous (e.g. *wt* sera and *wt* antigen) or heterologous (e.g. *wt* sera and ty-ad antigen) antigens. Individual points are shown with lines connecting paired sera. Paired t-tests were performed comparing the HAI titres of individual sera tested using the wt and ty-ad antigens. *** indicate p-value <0.001. ** indicate p-value <0.01.

**Fig. S4.** Representative IHC images from two turkeys infected with *wt* or ty-ad H7N9 viruses. Representative IHC images showing the presence of influenza A virus Nucleoprotein (NP) antigen on the indicated tissues collected from turkeys infected with *wt* or ty-ad H7N9 viruses. Tissues were collected from birds (*wt*, bird E or ty-ad, bird G) which were culled at 6 dpi. NP labelling is shown in brown.

**Fig. S5.** Representative images showing lectin staining for sialic acid receptor distribution in chicken and turkey tissues. Representative images showing the staining of α2,3 (Mal II) and α2,6 (SNA) sialic acid using lectins. Respective lectin staining of tissues collected from the nasal cavity of uninfected chickens and turkeys shown in brown.

**Fig. S6.** Representative IHC images from two ferrets infected with *wt* H7N9 viruses. Representative IHC images showing the presence of influenza A virus Nucleoprotein antigen in the indicated tissues collected from ferrets infected with the ty-ad H7N9 virus. Tissues were collected from ferrets which were culled at 6 dpi. NP labelling is shown in brown.

## REFERENCES

1. Gao R, Cao B, Hu Y, Feng Z, Wang D et al. Human infection with a novel avian-origin influenza A (H7N9) virus. N Engl J Med 2013;368(20):1888–1897.

2. World Health Organisation (WHO). 2021. Influenza at the human-animal interface summary and assessment, 21 May 2021. https://www.who.int/publications/m/item/influenza-at-the-human-animal-interface-summary-and-assessment-21may-2021 [accessed July 2023].

3. Food and Agriculture Organization of the United Nations (FAO). 2022. H7N9 situation update. https://www.fao.org/animal-health/situation-updates/avian-influenza-A(H7N9)-virus/en [accessed Nov 2023].

4. Chen Y, Liang W, Yang S, Wu N, Gao H et al. Human infections with the emerging avian influenza A H7N9 virus from wet market poultry: clinical analysis and characterisation of viral genome. Lancet 2013;381(9881):1916–1925.

5. Shi J, Deng G, Liu P, Zhou J, Guan L et al. Isolation and characterization of H7N9 viruses from live poultry markets — Implication of the source of current H7N9 infection in humans. Chinese Science Bulletin, journal article 2013;58(16):1857–1863.

6. Pantin-Jackwood MJ, Miller PJ, Spackman E, Swayne DE, Susta L et al. Role of poultry in the spread of novel H7N9 influenza virus in China. J Virol 2014;88(10):5381–5390.

7. Wang D, Yang L, Zhu W, Zhang Y, Zou S et al. Two Outbreak Sources of Influenza A (H7N9) Viruses Have Been Established in China. J Virol 2016;90(12):5561–5573.

8. Iuliano AD, Jang Y, Jones J, Davis CT, Wentworth DE et al. Increase in Human Infections with Avian Influenza A(H7N9) Virus During the Fifth Epidemic - China, October 2016-February 2017. MMWR Morb Mortal Wkly Rep 2017;66(9):254–255.

9. Zhu W, Dong J, Zhang Y, Yang L, Li X et al. A Gene Constellation in Avian Influenza A (H7N9) Viruses May Have Facilitated the Fifth Wave Outbreak in China. Cell reports 2018;23(3):909–917.

10. He D, Gu J, Gu M, Wu H, Li J et al. Genetic and antigenic diversity of H7N9 highly pathogenic avian influenza virus in China. Infection, Genetics and Evolution 2021;93:104993.

11. Yin X, Deng G, Zeng X, Cui P, Hou Y et al. Genetic and biological properties of H7N9 avian influenza viruses detected after application of the H7N9 poultry vaccine in China. PLOS Pathogens 2021;17(4):e1009561.

12. Wang Y, Lv Y, Niu X, Dong J, Feng P et al. L226Q Mutation on Influenza H7N9 Virus Hemagglutinin Increases Receptor-Binding Avidity and Leads to Biased Antigenicity Evaluation. J Virol 2020;94(20).

13. Peacock TP, Sealy JE, Harvey WT, Benton DJ, Reeve R et al. Genetic Determinants of Receptor-Binding Preference and Zoonotic Potential of H9N2 Avian Influenza Viruses. Journal of Virology 2021;95(5):e01651–01620.

14. Chang P, Sealy JE, Sadeyen J-R, Bhat S, Lukosaityte D et al. Immune Escape Adaptive Mutations in the H7N9 Avian Influenza Hemagglutinin Protein Increase Virus Replication Fitness and Decrease Pandemic Potential. Journal of Virology 2020;94(19):10.1128/jvi.00216-00220.

15. de Vries RP, Peng W, Grant OC, Thompson AJ, Zhu X et al. Three mutations switch H7N9 influenza to human-type receptor specificity. PLOS Pathogens 2017;13(6):e1006390.

16. Zhang H, Li X, Guo J, Li L, Chang C et al. The PB2 E627K mutation contributes to the high polymerase activity and enhanced replication of H7N9 influenza virus. J Gen Virol 2014;95(Pt 4):779–786.

17. Su S, Gu M, Liu D, Cui J, Gao GF, et al. Epidemiology, Evolution, and Pathogenesis of H7N9 Influenza Viruses in Five Epidemic Waves since 2013 in China. Trends Microbiol 2017;25(9):713–728.

18. Gilbert M, Golding N, Zhou H, Wint GRW, Robinson TP et al. Predicting the risk of avian influenza A H7N9 infection in live-poultry markets across Asia. *Nature Communications*, Article 2014;5:4116.

19. Lu Y, Landreth S, Gaba A, Hlasny M, Liu G et al. In Vivo Characterization of Avian Influenza A (H5N1) and (H7N9) Viruses Isolated from Canadian Travelers. Viruses 2019;11(2).

20. William T, Thevarajah B, Lee SF, Suleiman M, Jeffree MS et al. Avian influenza (H7N9) virus infection in Chinese tourist in Malaysia, 2014. Emerg Infect Dis 2015;21(1):142–145.

21. Shibata A, Okamatsu M, Sumiyoshi R, Matsuno K, Wang ZJ et al. Repeated detection of H7N9 avian influenza viruses in raw poultry meat illegally brought to Japan by international flight passengers. Virology 2018;524:10–17.

22. Bhat S, James J, Sadeyen JR, Mahmood S, Everest HJ et al. Co-infection of chickens with H9N2 and H7N9 avian influenza viruses leads to emergence of reassortant H9N9 virus with increased fitness for poultry and a zoonotic potential. J Virol 2022:jvi0185621.

23. James J, Bhat S, Walsh SK, Karunarathna TK, Sadeyen JR et al. The Origin of Internal Genes Contributes to the Replication and Transmission Fitness of H7N9 Avian Influenza Virus. J Virol 2022;96(22):e0129022.

24. Slomka MJ, Seekings AH, Mahmood S, Thomas S, Puranik A et al. Unexpected infection outcomes of China-origin H7N9 low pathogenicity avian influenza virus in turkeys. Sci Rep 2018;8(1):7322.

25. The Poultry Site. 2013. Global Poultry Trends 2013: Nearly One-fifth of Turkey Meat Exported. https://www.thepoultrysite.com/articles/global-poultry-trends-2013-nearly-onefifth-of-turkey-meat-exported [accessed Dec 2023].

26. Pillai SP, Pantin-Jackwood M, Yassine HM, Saif YM, Lee CW. The high susceptibility of turkeys to influenza viruses of different origins implies their importance as potential intermediate hosts. Avian Dis 2010;54(1 Suppl):522–526.

27. Spackman E, Gelb J, Preskenis LA, Ladman BS, Pope CR et al. The pathogenesis of low pathogenicity H7 avian influenza viruses in chickens, ducks and turkeys. Virology Journal 2010;7(1):331.

28. Malmberg JL, Miller M, Jennings-Gaines J, Allen SE. Mortality in Wild Turkeys (Meleagris gallopavo) Associated with Natural Infection with H5N1 Highly Pathogenic Avian Influenza Virus (HPAIV) Subclade 2.3.4.4. J Wildl Dis 2023;59(4):767–773.

29. Adlhoch C, Fusaro A, Gonzales JL, Kuiken T, Marangon S et al. Avian influenza overview March - June 2022. Efsa j 2022;20(8):e07415.

30. World Organisation for Animal Health (WOAH). Avian influenza (including infection with high pathogenicity avian influenza viruses). Manual of Diagnostic Tests and Vaccines for Terrestrial Animals. Paris, France: World Organisation for Animal Health (WOAH); 2023.

31. Reed LJ, Muench H. A simple method of estimating fifty percent endpoints. American Journal of Epidemiology 1938;27(3):493–497.

32. Seekings AH, Slomka MJ, Russell C, Howard WA, Choudhury B et al. Direct evidence of H7N7 avian influenza virus mutation from low to high virulence on a single poultry premises during an outbreak in free range chickens in the UK, 2008. Infection, Genetics and Evolution 2018;64:13–31.

33. Nagy A, Vostinakova V, Pirchanova Z, Cernikova L, Dirbakova Z et al. Development and evaluation of a one-step real-time RT-PCR assay for universal detection of influenza A viruses from avian and mammal species. Arch Virol 2010;155(5):665–673.

34. James J, Meyer SM, Hong HA, Dang C, Linh HTY et al. Intranasal Treatment of Ferrets with Inert Bacterial Spores Reduces Disease Caused by a Challenging H7N9 Avian Influenza Virus. Vaccines 2022;10(9):1559.

35. James J, Billington E, Warren CJ, De Sliva D, Di Genova C et al. Clade 2.3.4.4b H5N1 high pathogenicity avian influenza virus (HPAIV) from the 2021/22 epizootic is highly duck adapted and poorly adapted to chickens. J Gen Virol 2023;104(5).

36. Kumar S, Stecher G, Li M, Knyaz C, Tamura K. MEGA X: Molecular Evolutionary Genetics Analysis across Computing Platforms. Mol Biol Evol 2018;35(6):1547–1549.

37. Jiang W, Hou G, Li J, Peng C, Wang S et al. Prevalence of H7N9 subtype avian influenza viruses in poultry in China, 2013-2018. Transbound Emerg Dis 2019.

38. Blaurock C, Pfaff F, Scheibner D, Hoffmann B, Fusaro A et al. Evidence for Different Virulence Determinants and Host Response after Infection of Turkeys and Chickens with Highly Pathogenic H7N1 Avian Influenza Virus. Journal of Virology 2022;96(17):e00994–00922.

39. Kido H, Okumura Y, Takahashi E, Pan H-Y, Wang S et al. Role of host cellular proteases in the pathogenesis of influenza and influenza-induced multiple organ failure. Biochimica et Biophysica Acta (BBA) - Proteins and Proteomics 2012;1824(1):186–194.

40. Chaipan C, Kobasa D, Bertram S, Glowacka I, Steffen I et al. Proteolytic Activation of the 1918 Influenza Virus Hemagglutinin. Journal of Virology 2009;83(7):3200–3211.

41. Blaurock C, Scheibner D, Landmann M, Vallbracht M, Ulrich R et al. Non-basic amino acids in the hemagglutinin proteolytic cleavage site of a European H9N2 avian influenza virus modulate virulence in turkeys. Scientific Reports 2020;10(1):21226.

42. de Bruin ACM, Spronken MI, Bestebroer TM, Fouchier RAM, Richard M. Conserved Expression and Functionality of Furin between Chickens and Ducks as an Activating Protease of Highly Pathogenic Avian Influenza Virus Hemagglutinins. Microbiol Spectr 2023;11(2):e0460222.

43. Griffin DK, Robertson LB, Tempest HG, Vignal A, Fillon V et al. Whole genome comparative studies between chicken and turkey and their implications for avian genome evolution. BMC Genomics 2008;9(1):168.

44. Xiong X, Martin SR, Haire LF, Wharton SA, Daniels RS et al. Receptor binding by an H7N9 influenza virus from humans. Nature 2013;499(7459):496–499.

45. Shi Y, Zhang W, Wang F, Qi J, Wu Y et al. Structures and receptor binding of hemagglutinins from human-infecting H7N9 influenza viruses. Science 2013;342(6155):243–247.

46. Yang H, Carney PJ, Chang JC, Villanueva JM, Stevens J. Structural analysis of the hemagglutinin from the recent 2013 H7N9 influenza virus. J Virol 2013;87(22):12433–12446.

47. Kobayashi D, Hiono T, Ichii O, Nishihara S, Takase-Yoden S et al. Turkeys possess diverse Siaα2-3Gal glycans that facilitate their dual susceptibility to avian influenza viruses isolated from ducks and chickens. Virus Research 2022;315:198771.

48. Peacock TP, Benton DJ, James J, Sadeyen J-R, Chang P et al. Immune Escape Variants of H9N2 Influenza Viruses Containing Deletions at the Hemagglutinin Receptor Binding Site Retain Fitness *In Vivo* and Display Enhanced Zoonotic Characteristics. Journal of Virology 2017;91(14):e00218–00217.

49. Nilsson J, Eriksson P, Naguib MM, Jax E, Sihlbom C et al. Expression of influenza A virus glycan receptor candidates in mallard, chicken, and tufted duck. Glycobiology 2023.

50. Suzuki N, Abe T, Natsuka S. Structural analysis of N-glycans in chicken trachea and lung reveals potential receptors of chicken influenza viruses. Scientific Reports 2022;12(1):2081.

51. Singanayagam A, Zambon M, Barclay WS. Influenza Virus with Increased pH of Hemagglutinin Activation Has Improved Replication in Cell Culture but at the Cost of Infectivity in Human Airway Epithelium. Journal of Virology 2019;93(17):e00058–00019.

52. Zaraket H, Bridges OA, Russell CJ. The pH of activation of the hemagglutinin protein regulates H5N1 influenza virus replication and pathogenesis in mice. J Virol 2013;87(9):4826–4834.

53. Giannecchini S, Campitelli L, Calzoletti L, De Marco MA, Azzi A et al. Comparison of in vitro replication features of H7N3 influenza viruses from wild ducks and turkeys: potential implications for interspecies transmission. Journal of General Virology 2006;87(1):171–175.

54. Chang P, Sealy JE, Sadeyen JR, Iqbal M. Amino Acid Residue 217 in the Hemagglutinin Glycoprotein Is a Key Mediator of Avian Influenza H7N9 Virus Antigenicity. J Virol 2019;93(1).

55. James J, Billington E, Warren CJ, De Sliva D, Di Genova C et al. Clade 2.3.4.4b H5N1 high pathogenicity avian influenza virus (HPAIV) from the 2021/22 epizootic is highly duck adapted and poorly adapted to chickens. Journal of General Virology 2023;104(5).

