## Supplementary figures and images for "Evaluating the epizootic and zoonotic threat of an H7N9 low pathogenicity avian influenza virus (LPAIV) variant associated with enhanced pathogenicity in turkeys"

### Figure S1

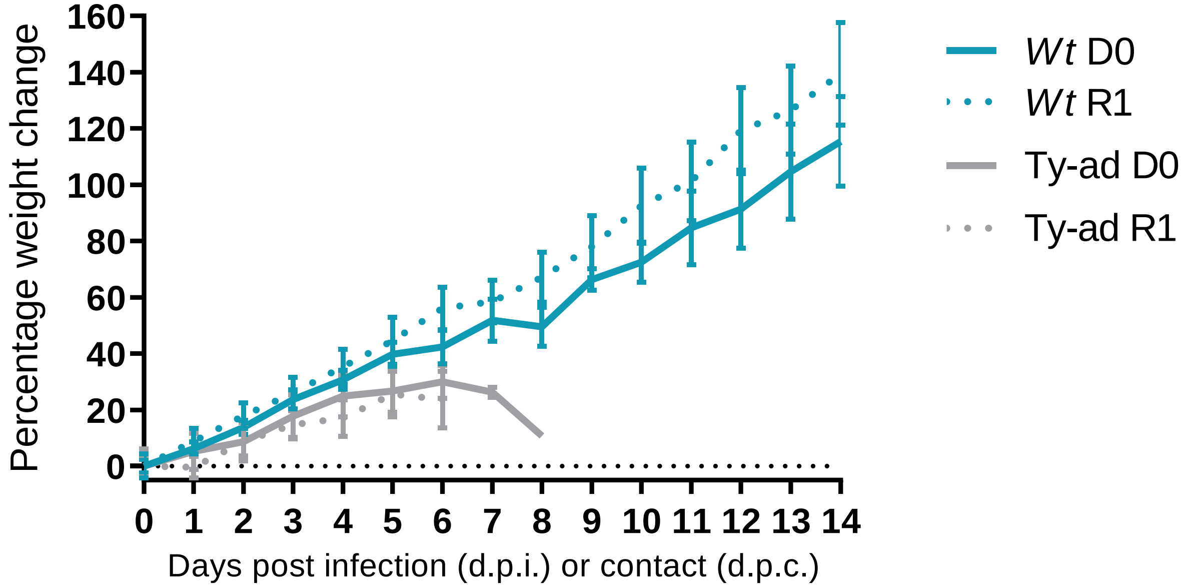

### Figure S2

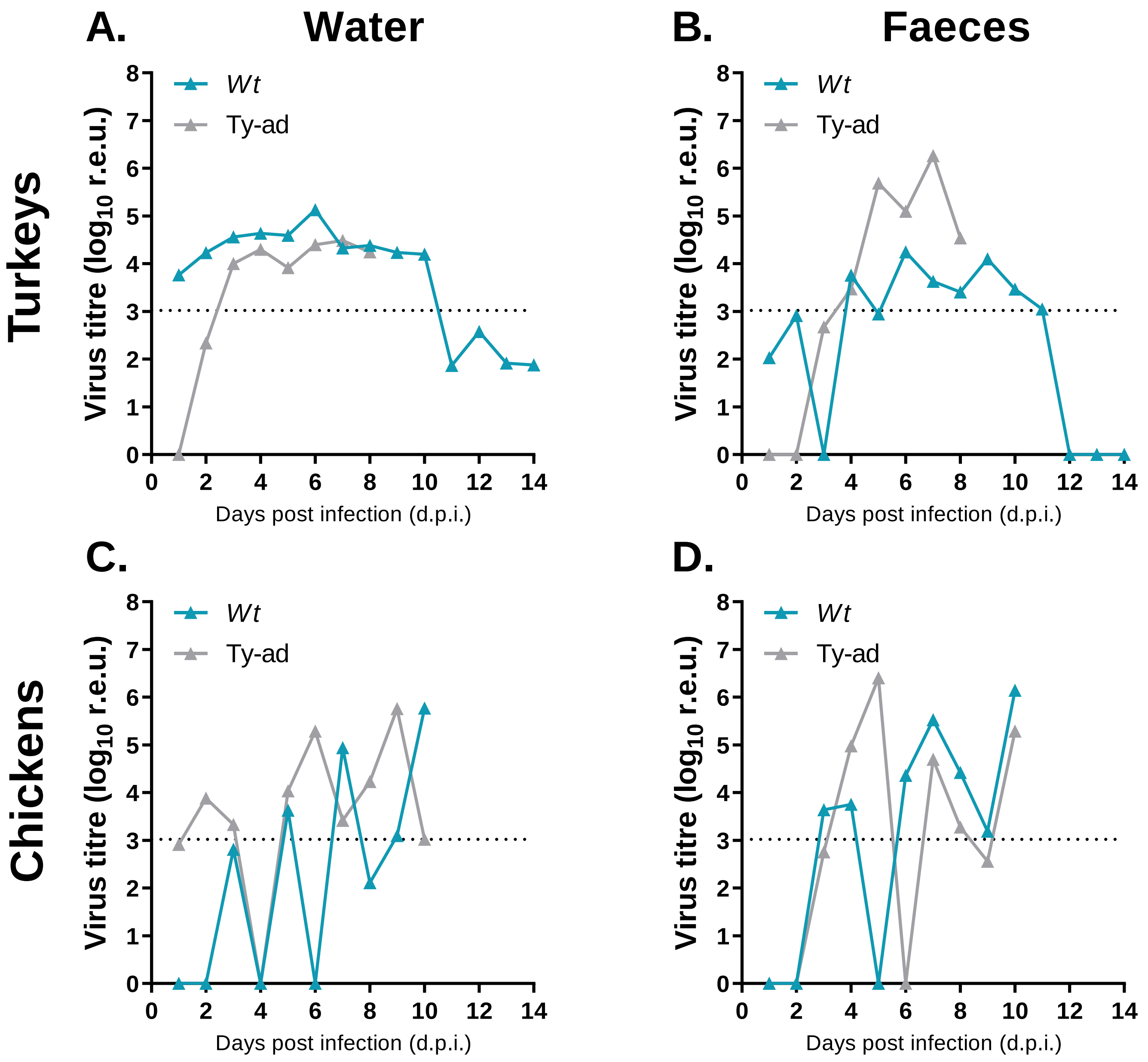

### Figure S3

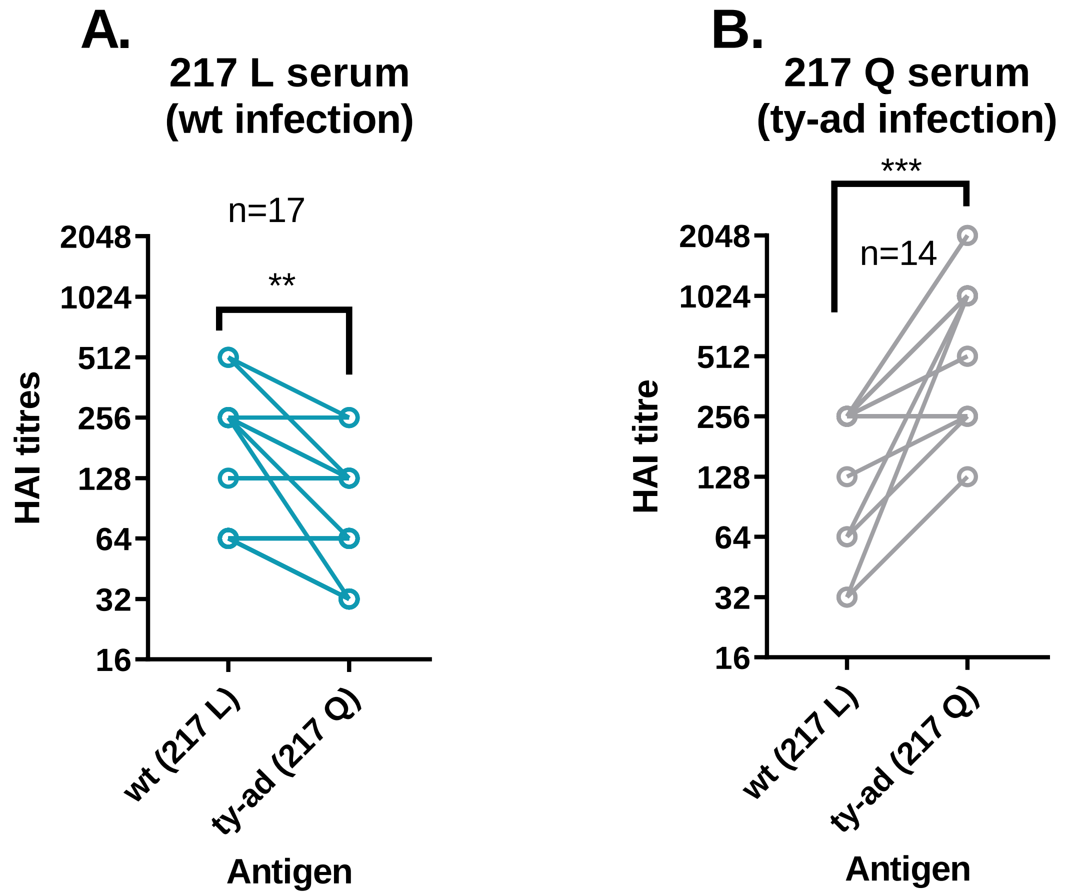

### Figure S4

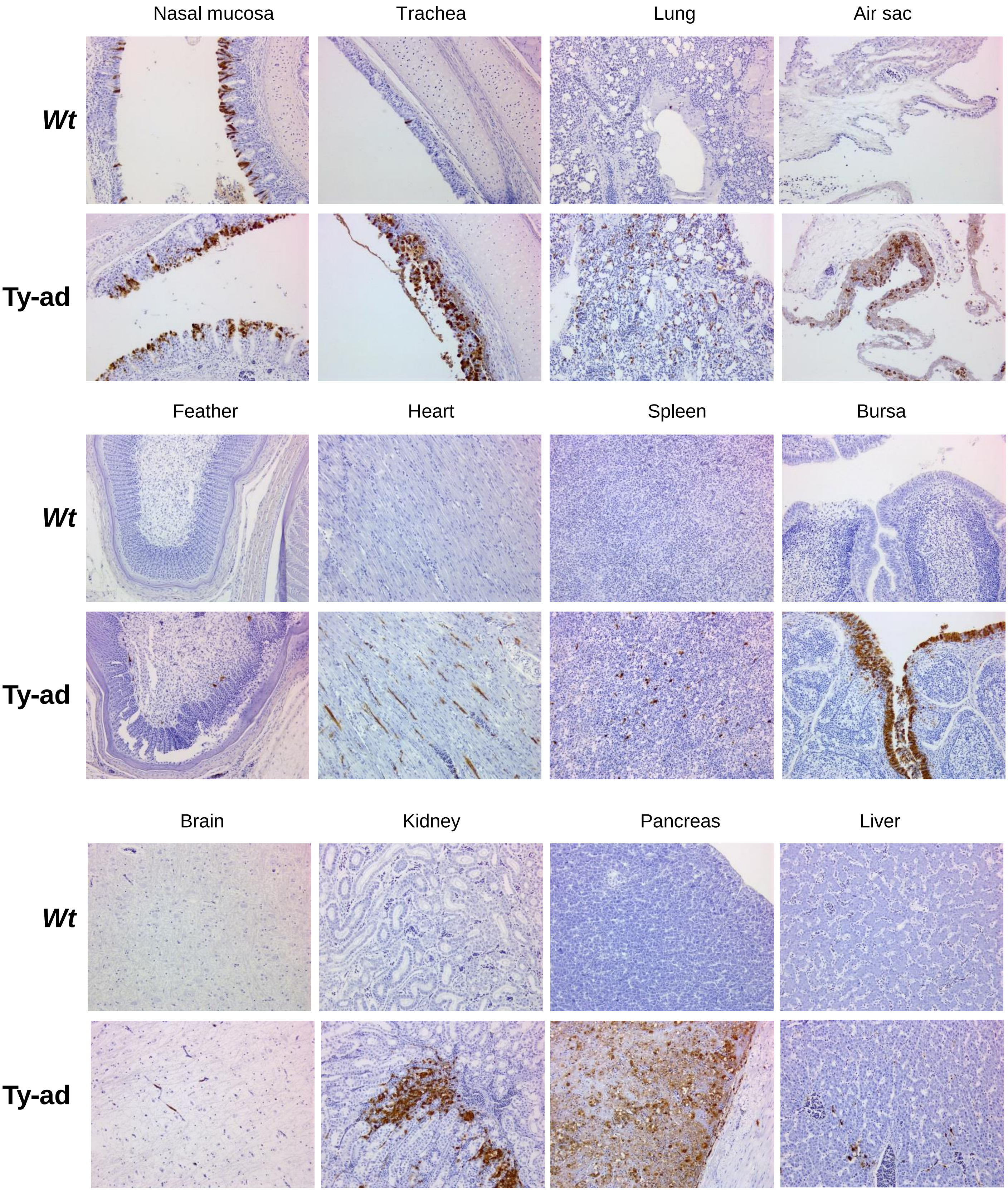

### Figure S5

#### Fig. S5

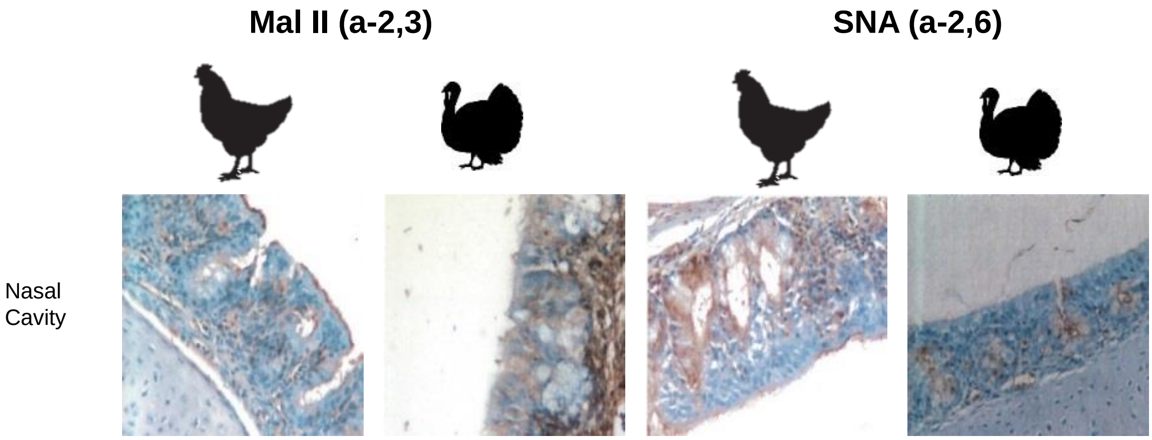

### Figure S6

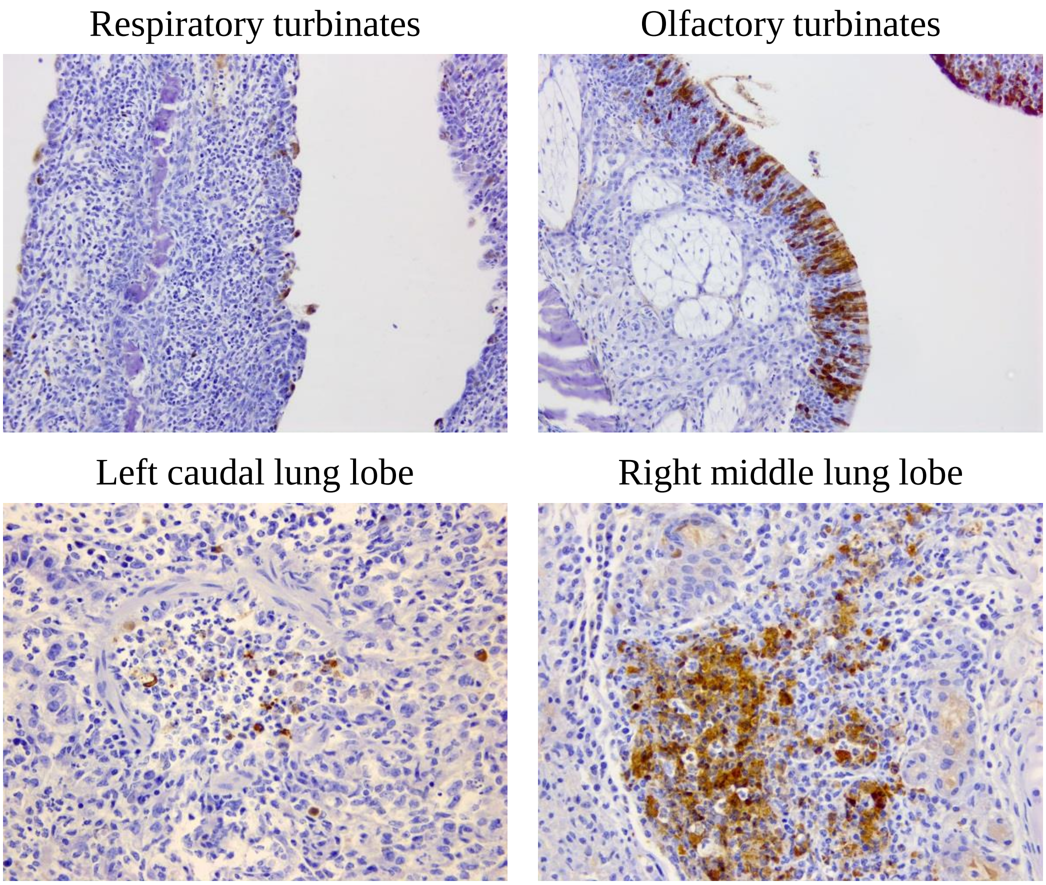
