## Supplementary material for "Evaluating the epizootic and zoonotic threat of an H7N9 low pathogenicity avian influenza virus (LPAIV) variant associated with enhanced pathogenicity in turkeys": Table S1

**Table S1.** Semi-quantitative evaluation of the influenza A type-specific NP antigen labelling by IHC. H7N9 LPAIV (wt and ty-ad) tropism was investigated in turkey tissues, following culls at 4 and 6 dpi when apparently healthy.

| | 4 DPI | | | | 6 DPI | | | |
|---|---|---|---|---|---|---|---|---|
| | Wt | | Ty-ad | | Wt | | Ty-ad | |
| **Bird ID** | **A** | **B** | **C** | **D** | **E** | **F** | **G** | **H** |
| **Heart** | - | - | ++ | +/- | - | - | + | ++ |
| **Skin** | - | - | + | - | - | - | +/- | + |
| **Feather follicles** | - | - | + | - | - | - | +/- | + |
| **Skeletal muscle** | - | - | + | - | - | - | +/- | + |
| **Spleen** | - | - | ++ | +/- | - | N/a | + | ++ |
| **Brain** | - | - | + | - | - | - | + | ++ |
| **Kidney** | - | - | + | - | - | - | + | ++ |
| **Ovary / Testes** | T - | T - | N/a | O - | O - | N/a | O + | O ++ |
| **Cecal Tonsil** | - | - | + | - | - | - | + | + |
| **Thymus** | - | - | + | - | - | - | +/- | +/- |
| **Bursa** | - | - | + | - | - | - | +++ | +++ |
| **Lung** | - | - | +++ | + | - | +/- | + | ++ |
| **Trachea** | - | - | +++ | + | - | + | ++ | +++ |
| **Air Sacs** | - | - | +++ | - | - | - | +++ | +++ |
| **Nasal cavity** | - | +++ | +++ | + | ++ | ++ | +++ | +++ |
| **Pancreas** | - | - | +++ | - | - | - | +++ | +++ |
| **Duodenum** | - | - | + | - | - | - | +/- | + |
| **Liver** | - | - | ++ | +/- | - | - | + | ++ |
| **Proventriculus** | - | - | + | - | - | - | +/- | + |
| **Jejunum** | - | - | + | - | - | - | +/- | + |
| **Colon** | - | - | + | - | - | - | +/- | + |
| **Caecum** | - | - | + | - | - | - | +/- | + |

O, ovary; T, testes; N/a, not available; -, absent; +/-, minimal; +, mild; ++, moderate; +++, diffuse
