## Supplementary material for "Evaluating the epizootic and zoonotic threat of an H7N9 low pathogenicity avian influenza virus (LPAIV) variant associated with enhanced pathogenicity in turkeys": Table S2

**Table S2.** Semi-quantitative evaluation of the influenza A NP antigen labelling by IHC in tissues collected from ferrets at 6 dpi following infection with the ty-ad H7N9 variant.

| Tissue | Detailed anatomical location | Ferret ID | |
|---|---|---|---|
| | | 1 | 2 |
| Turbinates | Nasal turbinate | + | + |
| | Olfactory turbinates | ++ | +/- |
| Glands | Salivary gland | - | - |
| Trachea | Cervical trachea | +/- | - |
| | Thoracic trachea | +/- | +/- |
| Lung lobe | Right cranial | - | - |
| | Right middle | +/- | ++ |
| | Right caudal | - | - |
| | Accessory | - | + |
| | Left cranial | - | - |
| | Left caudal | - | +/- |
| Lymph Nodes | Mandibular | - | - |
| | Retropharyngeal | - | - |

-, absent; +/-, minimal; + mild; ++, moderate; +++, diffuse.
